# DYRK3-Controlled Phase Separation Organizes the Early Secretory Pathway

**DOI:** 10.1101/2020.02.10.941757

**Authors:** Raffaella Gallo, Arpan Rai, Lucas Pelkmans

## Abstract

The dual-specificity kinase DYRK3 controls formation and dissolution of several intracellular condensates thereby regulating various cell physiological processes. Here we report that DYRK3 establishes a dynamic equilibrium between condensation and dissolution of proteins associated with membranous structures of the early secretory pathway to organize membrane traffic between the ER and the Golgi complex in mammalian cells. This depends on the peripheral membrane protein Sec16A, whose N-terminal disordered region forms DYRK3-controlled liquid-like condensates on the surface of the ER and co-phase separates with multiple ER exit site components and a subset of matrix proteins specifically associated with ERGIC and cis-Golgi. Our findings support a mechanism whereby multiple interacting and differentially regulated intracellular condensates create favorable environments for directional membrane traffic in eukaryotic cells.

## Introduction

The secretory pathway mediates the transport of most secretory and membrane proteins from the endoplasmic reticulum (ER) to the Golgi complex, and subsequently to the extracellular space, the cell surface or other membrane-bound organelles(*1–3*). At the ER, cargo is packed into coat protein complex II (COPII)-coated transport vesicles at specialized domains called ER exit sites (ERES). Extensive genetic, cell biological and biochemical work has characterized the machinery required for ERES and COPII vesicle formation and for establishing plasticity to adapt the size of transport carriers to the type of cargo(*3–10*). In this process, the peripheral membrane protein Sec16A plays an essential role by forming oligomers that may act as scaffolds for recruiting the components necessary for COPII vesicle formation(*5, 11–16*). How Sec16A achieves this remains largely unclear.

Recently, liquid-liquid phase separation (LLPS) has emerged as a general mechanism to locally concentrate multiple factors involved in complex biochemical processes(*17–20*). This typically involves weak multivalent interactions between proteins with intrinsically disordered regions (IDRs)(*21–24*). The resulting biomolecular condensates display rapid, liquid-like merging and can exchange components between the condensed and the dilute phase within seconds. Sec16A is also a highly disordered protein, with ∼75% of its total sequence predicted to be disordered in both *S. cerevisiae* and *H. sapiens*, as well as other ERES proteins including the transmembrane proteins TANGO1L (transport and Golgi organization protein 1) and cTAGE5 (cutaneous T cell lymphoma-associated antigen 5) (*25, 26*). Moreover, ERES components can exchange with a cytosolic pool within seconds, and individual ERES can rapidly merge with each other(*27–29*). Thus, the mechanism by which Sec16A acts as a central organizer of ERES may involve LLPS, which drives the formation of ER-associated biomolecular condensates that locally concentrate ERES proteins and COPII vesicle coat components. This behavior may also underlie the formation of Sec bodies in drosophila S2 cells, which are membraneless organelles formed by Sec16A during amino acid starvation(*30*). In addition, some COPII coat components partition into stress granules(*31*), which are membraneless organelles formed by LLPS during stress(*32, 33*), suggesting an affinity for biomolecular condensates.

As recently suggested(*34*), LLPS may also underly the formation of the Golgi matrix (*35*). This matrix consists of golgins, which are transmembrane or membrane-associated proteins with long coiled-coil regions in their cytosolic domains(*36, 37*). The cis-golgin GM130 has recently been shown to undergo LLPS *in vitro* and when overexpressed in cells(*38*). Similarly, TFG1 (Trk-fused gene 1), which interacts with Sec16A(*39*), forms a matrix at the interface between the ER and the ER-Golgi Intermediate Compartment (ERGIC) through oligomerization of C-terminal PQ (proline-glutamine)-rich IDRs(*40, 41*). Finally, the local clustering of synaptic vesicles in the synapse bares resemblance to that of COPII-coated vesicles at ERES(*40, 42–44*), which is achieved by a matrix-like synaptic condensate, driven by the IDR of the synaptic vesicle-associated protein Synapsin(*45*).

If LLPS organizes the early secretory pathway, it must be actively controlled to support the dynamic properties of vesicular traffic and the directional flux of cargo. In agreement with this, ATP depletion causes the appearance of a fibrous dense matrix between the cis-Golgi and the ER (*46, 47*), indicating that its condensation properties are actively controlled (*28, 48–50*). A key mechanism of controlling the condensation properties of intracellular condensates is by means of reversible phosphorylation. Dual-specificity tyrosine-regulated kinase (DYRK) family members have emerged as central regulators of LLPS through phosphorylating their substrates in IDRs, mediating the dissolution of stress granules, P granules, pericentriolar satellites, nuclear speckles, and PML nuclear bodies(*32, 51–53*). Interestingly, in the original RNAi screen that revealed DYRK3 as a host factor involved in virus infection, it was reported that its depletion affected the morphology of membranous structures carrying retrograde transport cargo(*54*). This was corroborated in a more recent screen, showing that knockdown of DYRK3 affects the secretory pathway(*55*). Here, we identify DYRK3 as a critical component in the machinery controlling the liquid-liquid phase behavior of the early secretory pathway by direct regulation of the condensation behavior of Sec16A, the most upstream component of ERES.

### DYRK3 interacts with COPII proteins and its inhibition condenses early secretory pathway components

Using quantitative proteomics in HEK293T cells, we previously identified 251 proteins that specifically interact with DYRK3(*52*), including Sec16A and the COPII coat components Sec13, Sec23A/B, and Sec24A/B (Fig. 1A, Suppl. Fig 1A). The Sec proteins were amongst the top-15 factors identified in the interactome(*52*) raising the possibility that DYRK3 engages with the components of the early secretory pathway in a specific and functionally relevant fashion. To test this idea, we expressed GFP-tagged DYRK3 below endogenous levels, and occasionally observed DYRK3 in structures that also contained endogenous Sec16A and Sec13, which mark ERES (Suppl. Fig. 1B). The localization of DYRK3 to ERES became more prominent upon acute (1h) inhibition of DYRK3 using the small compound inhibitor GSK-626616 (Fig. 1B), which resulted in an increase in ERES size (Suppl. Fig. 1C). This behavior is reminiscent of other DYRK3-controlled intracellular condensates, which dissolve less efficiently when the kinase activity of DYRK3 is inhibited(*32, 52*).

**Figure 1.**
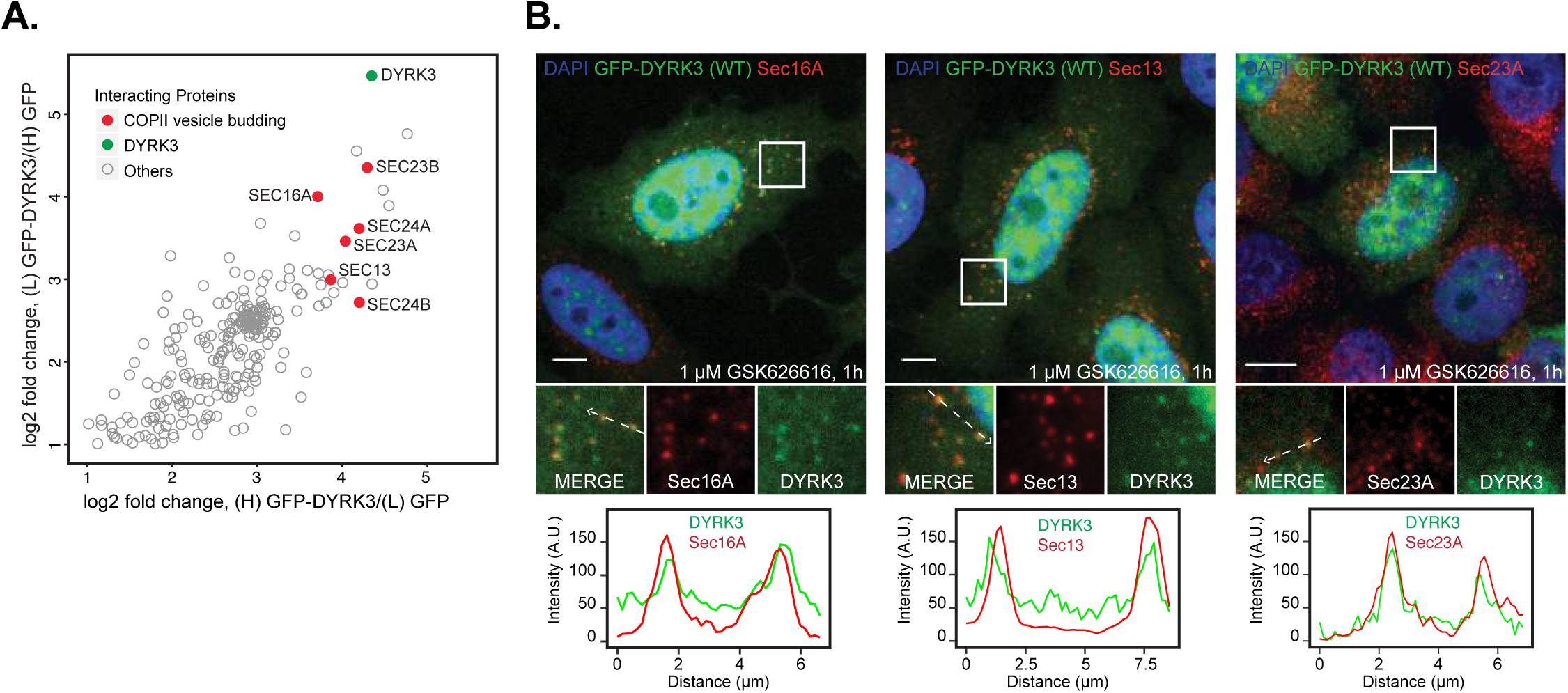
DYRK3 interacts with COPII machinery and localize to ER exit sites. A. Scatter plot shows enrichment of DYRK3 specific interactions compared to GFP control. Proteins with normalized log2 ratios higher than the cut-off value of 1, in both the forward and reverse pull-downs, are considered specific interaction partners. Proteins associated with COPII vesicle budding are color-coded (*52*). B. Upper Panel: Immunofluorescence images of cells (HeLa Flp-In T-Rex) show co-localization of GFP-tagged wildtype DYRK3 (GFP-DYRK3 (WT)) with COPII associated proteins in the presence of GSK626616 (1µM, 1h). Lower Panel: The fluorescent line profile of GFP-DYRK3 and ER exit site (ERES) markers along the arrows in the inset. GFP-DYRK3 (WT) expression was induced in HeLa Flp-In T-Rex cells by adding doxycycline (500ng/mL, 4hrs). Images are representative of at least three independent experiments. All scale bars: 10 µm.

Consistently, prolonged treatment of two mammalian cell lines (HeLa and Cos7) with GSK-626616 (3h), caused a strong size increase of Sec16A- and Sec13-positive structures and lead to the accumulation of these structures in the perinuclear region (Fig 2A, Suppl. Fig. 2A,B). The aberrant Sec16A- and Sec13-positive structures also co-clustered with markers of the ER-Golgi intermediate compartment (ERGIC53), as well as with the cis-Golgi complex component GM130 (Fig 2A, Suppl. Fig. 2A,B). Inhibition of DYRK3 therefore resulted in profound re-arrangements of all organelles involved in the early secretory pathway preventing the typical juxtapositioning of Sec16A- and ERGIC53, and severely perturbing the extended ribbon-like organization of the Golgi complex (Fig 2A, Suppl. Fig. 2A, B). The ER itself appeared largely unperturbed (Fig. 2B, Suppl. Fig 2C). Time-lapse imaging of GFP-Sec16A-expressing cells revealed that peripheral ERES structures disappeared upon addition of GSK-626616, and gradually accumulated in enlarged perinuclear structures (Fig. 2C, D). Importantly, these structures did not co-stain for G3BP1 and ubiquitin, indicating that they were not stress granules or aggresome-like accumulations of misfolded proteins (Suppl. Fig. 2D-F). The perinuclear accumulation of Sec16A was completely reversed within 1h after removing the inhibitor (Suppl. Fig. 2G).

**Figure 2.**
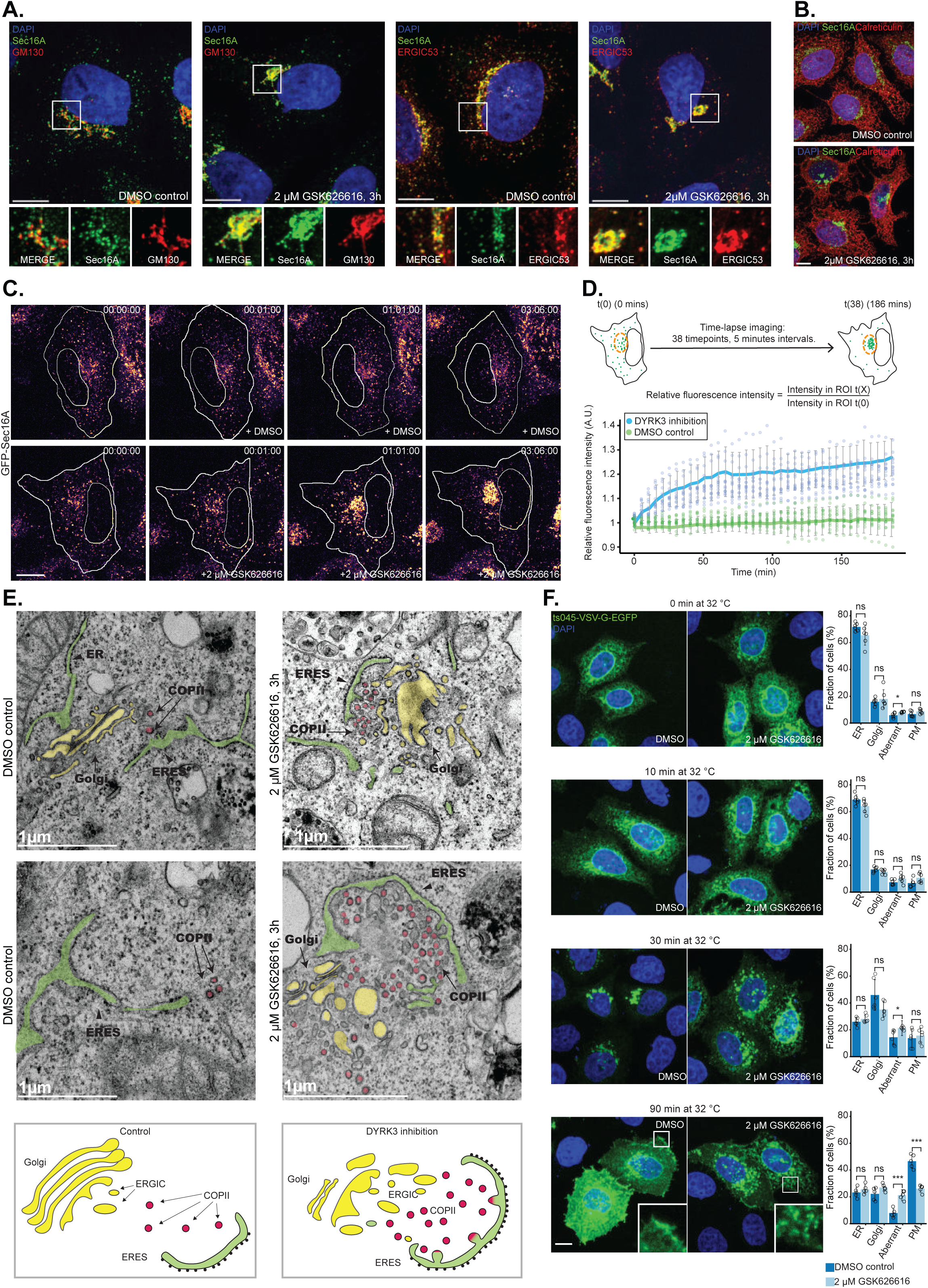
DYRK3 inhibition perturbs organization of the ER-Golgi interface and inhibits secretory trafficking. A. Immunofluorescence images of cells (HeLa) stained with ERES marker (Sec16A), Golgi marker (GM130) and ERGIC marker (ERGIC53) show perinuclear clustering of these components upon GSK626616 treatment (2µM, 3h). B. Immunofluorescence images of cells (HeLa) stained with ER marker (Calreticulin) and ERES marker (Sec16A) show ER network is unperturbed upon GSK626616 treatment (2µM, 3h). C. Time lapse images of cells (HeLa) overexpressing full-length GFP-Sec16A show perinuclear accumulation of Sec16A upon GSK626616 treatment (2µM, indicated times). D. Quantification of the relative change in fluorescence intensity in the perinuclear region (ROI) over time (Fig. 2C) upon GSK-626616 treatment (2µM), compared to DMSO control. Data are mean ± s.d. (n=15 cells per condition, 2 independent experiments). Schematic depicting methodology used for quantification is shown above. E. Electron microscopy images of cells (HeLa) show accumulation of vesicles, vesicle budding intermediates and swollen Golgi at ER-Golgi interface upon GSK626616 treatment (2µM, 3h) (n=10 cells per conditions, 2 independent experiments). ER, ERES, Golgi apparatus and likely COPII vesicles are marked in the images. Schematic depicting observed changes at ER-Golgi interface upon GSK626616 treatment is shown. F. (Left) Immunofluorescence images of cells (HeLa) show transport of ts045-VSV-G-GFP from the endoplasmic reticulum (ER) to the plasma membrane (PM) through the Golgi apparatus (Golgi) upon temperature shift from 40°C to 32°C, is perturbed in the presence of GSK626616 (2µM). The indicated compounds were added an hour before the temperature shift. (Right) Quantification of the fraction of cells in the four classes (classified by SVM training) at indicated time points. Time points represent time after shift to permissive temperature (32°C). Data are mean ± s.d (Data shown is from 3 independent experiments). Statistical analysis was performed across the four classes comparing DMSO and GSK626616 treatment using Student’s t-test. * - P < 0.05, *** - P < 0.001, ns - not significant. Images are representative of at least three independent experiments. All scale bars (except Fig. 2E): 10 µm.

We then used electron microscopy to look at the ultrastructural organization of those perinuclear accumulations. Control cells showed regions in which ER cisternae, a few vesicular structures resembling COPII-coated vesicles, and a typically stacked Golgi complex are in close proximity to each other (Fig. 2E). In GSK-626616-treated cells, such regions contained an unusually large amount of vesicular structures resembling COPII-coated vesicles (Fig. 2E) that had accumulated between swollen ERGIC- and cis-Golgi-like intermediates and an enlarged ERES containing multiple vesicle budding intermediates.

To determine whether the reorganization of the ER-to-Golgi interface induced by the inhibition of DYRK3 perturbs secretory trafficking, we overexpressed in cells a temperature-sensitive folding mutant of Vesicular Stomatitis Virus G protein fused to GFP (ts045-VSV-G-GFP). Ts045-VSV-G-GFP misfolds and is retained in the ER when cells are incubated at the non-permissive temperature (40°C) but gets rapidly folded and exported upon shift to a permissive temperature (32°C)(*56*). In control cells, ts045-VSV-G-GFP behaved as expected, accumulating in a perinuclear ribbon-like Golgi complex 30 min after shifting the cells to permissive temperature and readily reaching the cell surface after 90 min (Fig. 2F). In inhibitor-treated cells, shifting to the permissive temperature resulted in ts045-VSV-G-GFP accumulating in the aberrant structures as describe above, and was unable to reach the plasma membrane even after 90 min (Fig. 2F). However, a similar fraction of ts045-VSV-G-GFP became Endonuclease H (EndoH)-resistant in both control and GSK-626616-treated cells, indicating that cargo becomes exposed to terminal N-linked glycosylation enzymes normally resident in the cis-Golgi (Suppl. Fig. 3). Thus, DYRK3 inhibition has severe effects on organization and trafficking in the early secretory pathway.

### Overexpression of DYRK3 dissolves ER exit sites and collapses the Golgi complex

If DYRK3 inhibition results in an over-condensation of ERES, ERGIC, and cis-Golgi, then, in analogy to its effect on other intracellular condensates(*52*), DYRK3 overexpression would lead to its dissolution. To explore this hypothesis, we transiently transfected GFP-DYRK3 in cells and stained against endogenous COPII proteins. Upon GFP-DYRK3 overexpression, the typical spot-like Sec16A and Sec13 distribution disappeared, resulting in a more diffuse cytoplasmic signal (Fig. 3A, B). This dissolution was switch-like, occurring only above a critical DYRK3 concentration (Fig. 3C). It also resulted in a complete redistribution of the transmembrane protein TANGO1L, which depends on Sec16A for its localization at ERES(*57*), across the ER, co-localizing with calreticulin (CRT). (Fig. 3D), and led to a striking re-localization of the Golgi matrix protein GM130 into a condensed structure in the center of the cell (Fig. 3E). Overexpression of a kinase-dead point mutant of DYRK3 (K218M) did not have these effects, while treating DYRK3-overexpressing cells with GSK-626616 reversed the effect (Fig. 3A, B; Suppl. Fig. 4A, B), resulting in the entrapment of GFP-DYRK3 in TANGO1L-positive ERES (Suppl. Fig. 4C). As a consequence of ERES loss, functional secretory trafficking was blocked, with ts045-VSV-G-mCherry being trapped in the ER upon DYRK3 overexpression, even after 90 min at the permissive temperature, where it remained EndoH-sensitive (Fig. 3D, E). Among all examined DYRK family members, as well as CDK1/CyclinB, only its closest family member DYRK2 was able to partly recapitulate the DYRK3 overexpression effects, but at higher expression levels (Suppl. Fig. 4D). Thus, through its kinase activity and ability to partition into ERES, DYRK3 is critically involved in controlling the condensation threshold of ERES proteins, thereby determining ERES nucleation and regulating ER export.

**Figure 3.**
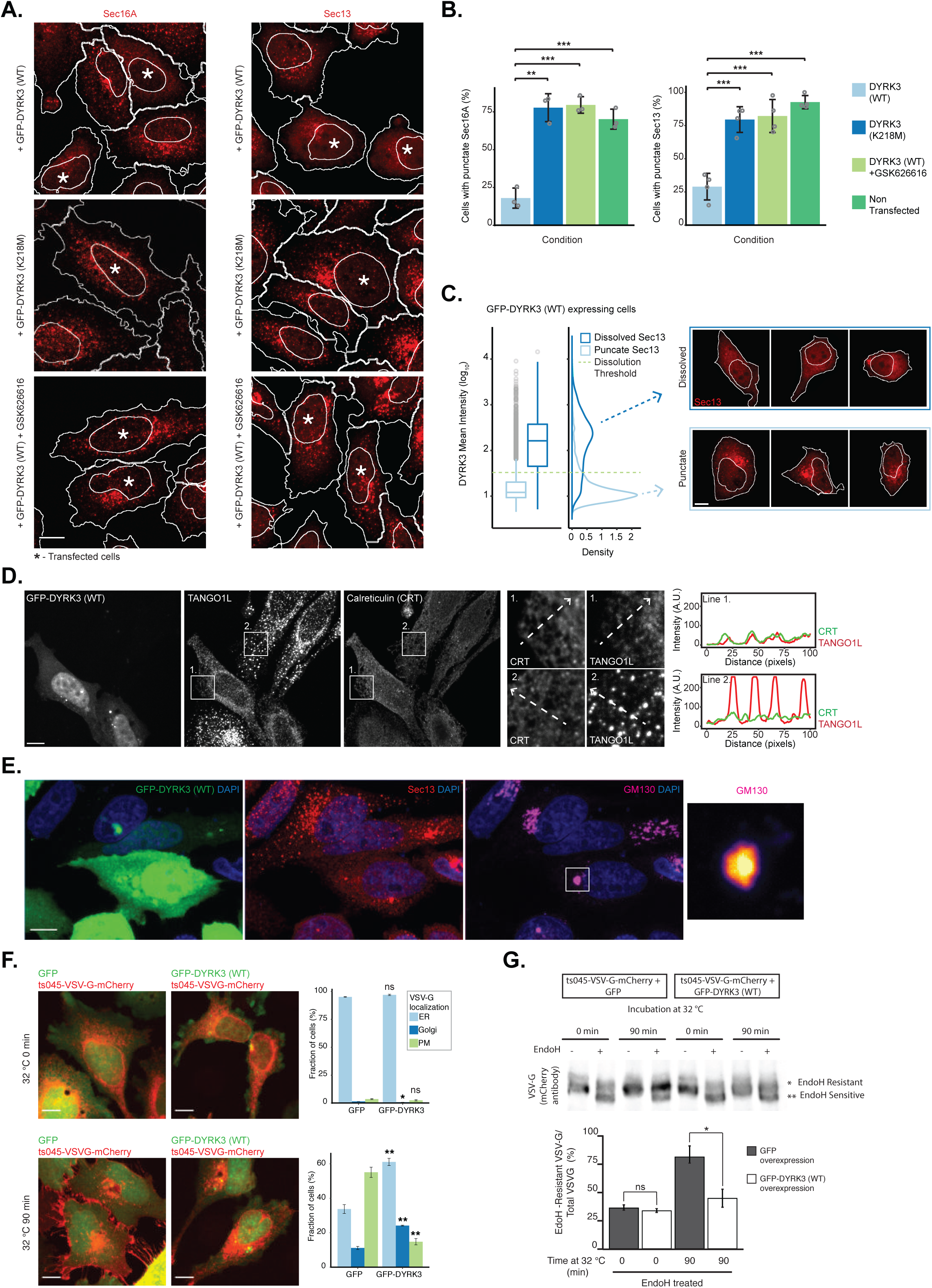
DYRK3 overexpression dissolves ER exit sites and inhibits secretory trafficking. A. Immunofluorescence images of cells (HeLa) show dissolution of punctate ERES markers (Sec16A and Sec13) upon overexpression of wild-type DYRK3 (GFP-DYRK3 (WT)), but not kinase-dead DYRK3 ((GFP-DYRK3 (K218M)). Dissolution of ERES markers by wild-type DYRK3 is reversed upon addition of the inhibitor, GSK-626616 (1µM, 1h). Asterisks indicate transfected cells. GSK626616 instead of inhibitor. B. Quantification of the dissolution of ERES puncta (Sec16A and Sec13) by DYRK3. Data are mean ± s.d (Data shown is from 3 independent experiments). Statistical analysis was performed using Student’s t-test. ** P < 0.01,*** P <0.001 C. Quantification of the dissolution of Sec13 puncta as a function of DYRK3 expression level shows DYRK3 regulates ERES dissolution in a concentration-dependent manner. Cells containing a punctate or dissolved Sec13 staining in HeLa cells were classified by SVM training. Representative images of dissolved and punctate Sec13 staining are shown. Data shown is from 3 technical replicates. D. Immunofluorescence images of cells (HeLa) show redistribution of punctate TANGO1L across the ER network (Calreticulin (CRT)) upon overexpression of GFP-DYRK3 (WT). The fluorescent line profile of TANGO1L and CRT along the arrows in the inset is shown. E. Immunofluorescence images show cells (HeLa) with dissolved ERES (Sec13) upon overexpression of GFP-DYRK3 (WT), have a condensed Golgi structure (GM130) in the perinuclear region. F. Immunofluorescence images of cells (HeLa) show transport of VSVG (ts045-VSV-G-mCherry) from the endoplasmic reticulum (ER) to the plasma membrane (PM) through the Golgi apparatus (Golgi) upon temperature shift from 40°C to 32°C, is blocked upon overexpression of GFP-DYRK3 (WT). Quantification of the fraction of cells in the three classes (classified by SVM training) at indicated time points. Time points represent time after shift to permissive temperature (32°C). Data are mean ± s.d. (Data shown is from 3 independent experiments). Statistical analysis was performed across the three classes comparing overexpression of GFP-DYRK3 (WT) with GFP using Student’s t-test. * - P < 0.05, ** P < 0.01, ns - not significant. G. Representative western blot shows evaluation of glycosylation of VSVG (ts045-VSV-G-mCherry) by endoglycosidase H (Endo H) sensitivity assay at the permissive temperature (32°C) for indicated time points. VSVG remained EndoH sensitive after 90 min at the permissive temperature (32°C) upon GFP-DYRK3 (WT) overexpression. EndoH sensitive and resistant form of VSVG is labeled in the blot. Quantification of fraction of EndoH resistant VSVG form after indicated time points at the permissive temperature (32°C). Data are mean ± s.d. (Data shown is from 3 independent experiments). Statistical analysis was performed using Student’s t-test. * - P < 0.05, ns - not significant. Images are representative of at least three independent experiments. All scale bars: 10 µm.

### Sec16A-IDR forms liquid-like condensates that recruit early secretory pathway proteins and enwrap ER

To further investigate how the condensate properties of ERES depend on DYRK3 activity, we developed an experimental system that easily allows the visualization of this process in living cells. Using a set of Sec16A truncation mutants in a structure-function analysis we found that Sec16A1-1,227 results in the appearance of spherical structures in cells that are different from the typical ERES localization of the full-length protein (Fig. 4A, B and Suppl. Fig. 5A). This region of Sec16A (which we term Sec16A-IDR) is intrinsically disordered and comprises slightly more than half of its full-length sequence, including the N-terminal part and the previously reported ER localization domain (ELD)(*13, 58*). In time-lapse imaging, these structures are reminiscent of condensates, whose content has unmixed from the surrounding cytoplasm and that undergo liquid-like merging (Fig. 4C). Using photobleaching, we observed that the exchange of Sec16A-IDR molecules between the condensed droplet and the cytoplasm occurred within seconds (Fig. 4D). Importantly, the Sec16A-IDR condensates did not recruit the stress granule marker G3BP1, indicating that they are a different type of condensate. (Suppl. Fig. 5B). To characterize their composition, we stained for several endogenously expressed early secretory pathway components and found that most of them localized to the Sec16A-IDR condensates. Besides endogenous full length Sec16A, we detected the COPII coat components Sec13 and Sec23A, as well as TFG1, a protein enriched at the ER-Golgi interface(*40, 41*), GM130, a cis-Golgi golgin, and p115, which interacts with GM130 to form receptors for incoming transport vesicles from the ER, within the condensates(*59, 60*) (Fig 4E). Interestingly, also the transmembrane proteins Sec12 and TANGO1L, as well as co-expressed mCherry-cTAGE5, were sequestered into some of the condensates (Fig. 4E, Suppl. Fig. 5C). Possibly, Sec16A-IDR condensates that are in close proximity to the ER can act as a sink for these transmembrane proteins. Consistently, staining against endogenous calreticulin showed that some of these condensates (Fig. 4F ROI I, II, II) but not all (Fig. 4F ROI IV) were in close proximity to the ER membranes. Furthermore, correlative light-electron microscopy (CLEM) showed that the Sec16A-IDR condensates are amorphous round accumulations of electron-dense material inside the cytoplasm (Fig. 4G). When they were in close proximity to the ER, they had a profound effect on local ER morphology. Occasionally, condensates appeared to have formed extensive contacts with the surface of the ER, resulting in the exclusion of ribosomes, reminiscent of ERES being ribosome-free(*47, 61*). These ER cisternae appeared dilated accumulating large amounts of material in the lumen. Occasionally, we observed a Sec16A-IDR condensate to have engulfed multiple dilated ER cisternae (Fig. 4G). Consistent with the profound effects of the Sec16A-IDR condensates on the localization of early secretory pathway components and ER morphology, we found that cells overexpressing GFP-Sec16A-IDR were strongly compromised in their ability to export secretory cargo from the ER, when compared to cells overexpressing full length Sec16A or GFP alone (Fig. 4H).

**Figure 4.**
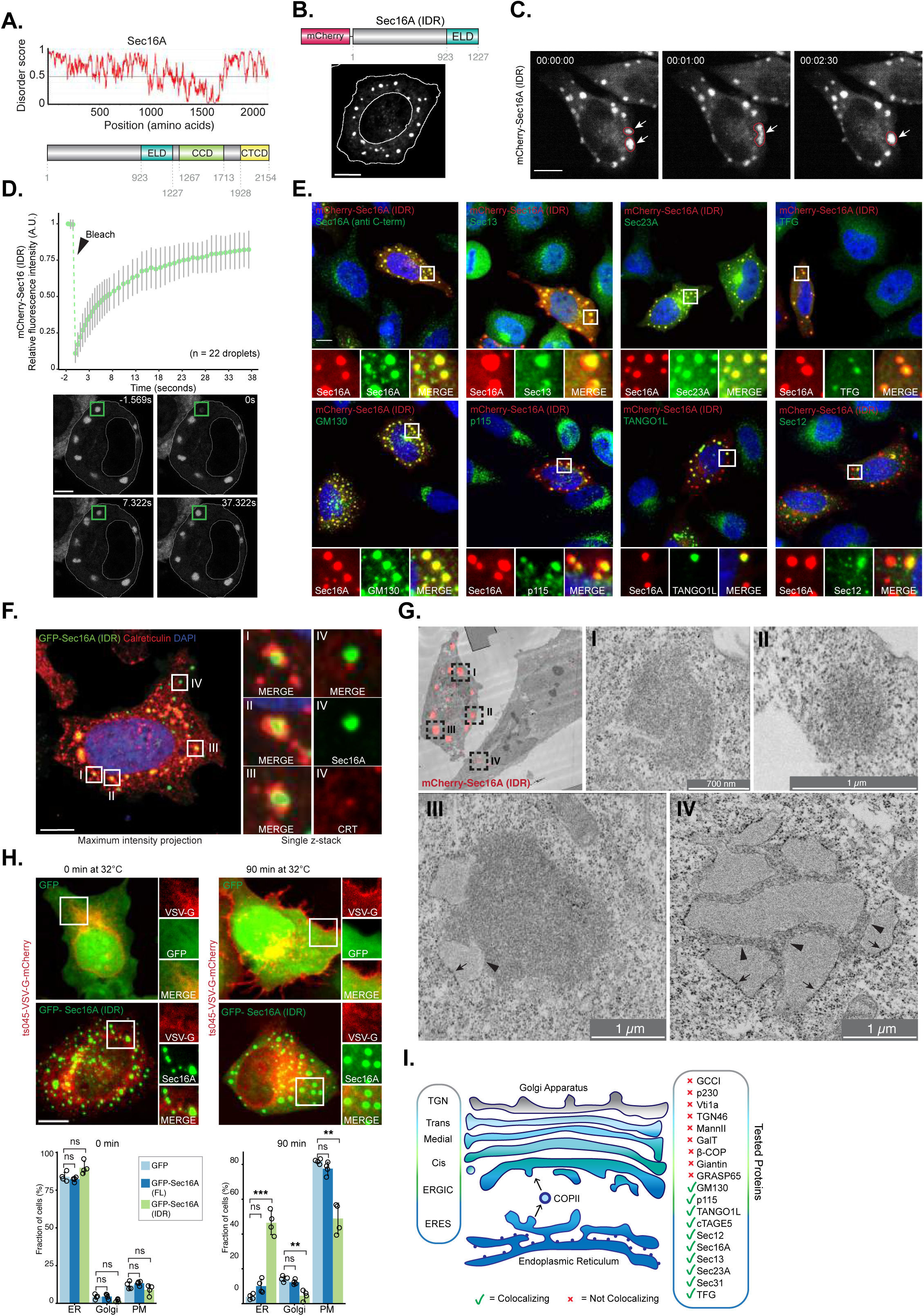
Sec16-IDR forms liquid-like condensates that recruit early secretory pathway proteins. A. Intrinsic disorder analysis of Sec16A using IUPRED prediction tool is shown. Schematic depicting the functional domains in full-length Sec16A. ELD: ER localizing domain; CCD: Central conserved domain; CTCD: C-terminal conserved domain B. Immunofluorescence image of a cell (HeLa) shows condensate formation by over-expressed mCherry-Sec16A (IDR). C. Time-lapse images of a cell (HeLa) (single Z-stack) show fusion of mCherry-Sec16A (IDR) condensates (shown with arrowheads). D. Fluorescence recovery after photobleaching (FRAP) trajectories of mCherry-Sec16A (IDR) condensates in cells (HeLa). Images show FRAP recovery. Data are mean ± s.d (n=12 cells, 3 independent experiments). E. Immunofluorescence images of cells (HeLa) show recruitment of endogenous ERES proteins (Sec16A, Sec23A, Sec12, TANGO1L), ER-ERGIC associated matrix protein (TFG) and cis-Golgi associated proteins (GM130 and p115) to mCherry-Sec16A (IDR) condensates. Endogenous Sec16A was stained with an antibody targeting the C-terminal end of the protein. F. Immunofluorescence image of a cell (HeLa) shows ER (Calreticulin (CRT)) in some cases wrapped around the condensates formed by GFP-Sec16A (IDR). G. Correlative light-electron microscopy images of cells (HeLa) show mCherry-Sec16A (IDR) condensates are membraneless compartments, some of which are in close proximity to the ER membrane. Arrowheads and arrow indicate ribosome-free and ribosome-attached surfaces of the ER (n=10 cells, 2 independent experiments). H. Upper Panel: Immunofluorescence images of cells (HeLa) show transport of VSVG (ts045-VSV-G-mCherry) from the endoplasmic reticulum (ER) to the plasma membrane (PM) through the Golgi apparatus (Golgi) upon temperature shift from 40°C to 32°C, is blocked upon overexpression of GFP-Sec16A (IDR). Lower Panel: Quantification of the fraction of cells in the three classes (classified by SVM training) at indicated time points. Data are mean ± s.d (Data shown is from 2 technical replicates per experiment, 2 independent experiments). Statistical analysis was performed across the three classes comparing overexpression of GFP-Sec16A full-length (FL) and GFP-Sec16A (IDR) with GFP using Student’s t-test. * - P < 0.05, ** P < 0.01, *** - P < 0.001, ns - not significant. I. Schematic representation of proteins of the ER and Golgi compartments that were tested for recruitment (colocalize) to condensates formed by Sec16A (IDR). Images are representative of at least three independent experiments. All scale bars (except 4G): 10 µm.

Although the Sec16A-IDR condensates recruited multiple proteins involved in early secretory transport between the ER and the cis-Golgi, they did not recruit the membrane-associated or transmembrane proteins of the cis-, medial- or trans-Golgi network (TGN) GRASP65, Giantin, p230 (Golgin-245), GCC1, Vti1a, and TGN46, or the luminal Golgi-resident proteins 1,4-galactosyltransferase (GalT) and a-mannosidase II (MannII) (Suppl. Fig. 5D). The Golgi complex often appeared fragmented under these conditions, and Golgi-resident enzymes accumulated in the ER, indicating that secretory membrane traffic is compromised. Finally, we did not detect components of the COPI vesicle coat (betaCOP) in the Sec16-IDR condensates (Suppl. Fig 5D).

Thus, the IDR of Sec16A forms, when overexpressed, liquid-like condensates inside cells that induce the co-condensation of multiple proteins that are involved in ERES formation and ER-to-Golgi membrane traffic. These condensates can recruit cytosolic (Sec13, Sec23, Sec31A), membrane-associated (TFG, GM130, and p115), and transmembrane proteins with long cytosolic domains (Sec12, TANGO1L, and cTAGE5), but importantly, exclude proteins that are associated with the medial-Golgi and TGN (Fig. 4I).

### DYRK3 modulates Sec16A-IDR condensates through phosphorylation of Sec16A

To test whether Sec16A-IDR condensates are under control of DYRK3, we transfected mCherry-Sec16A1-1,227 in cells containing inducible, GFP-tagged DYRK3 and performed timelapse imaging to follow the phase transition behavior of the condensates after GFP-DYRK3 induction. At low expression levels, we observed that GFP-DYRK3 partitioned into Sec16A-IDR condensates causing their rapid dissolution as its expression levels increased (Fig. 5A). This is reminiscent of the effect DYRK3 has on SRRM1, a phase-separating protein of nuclear speckles, that crosses the phase boundary upon overexpression of DYRK3(*52*). Sec16A-IDR condensate dissolution was readily reversed by adding GSK-626616, resulting in the reformation of condensates within 5 min (Fig. 5A). Testing the effect of the serine/threonine phosphatase inhibitors Calyculin A and Okadaic acid on cells containing Sec16-IDR condensates showed a potent, opposite effect. Sec16-IDR condensates were rapidly dissolved after 14 min of adding the inhibitor, which was reversed within 14 min of inhibitor washout (Fig. 5B, Suppl. Fig. 6A). This indicates that Sec16A-IDR condensates are continuously modulated by DYRK3, and their condensation threshold is determined by a dynamic equilibrium between DYRK3 and one, or multiple, serine/threonine phosphatases.

**Figure 5.**
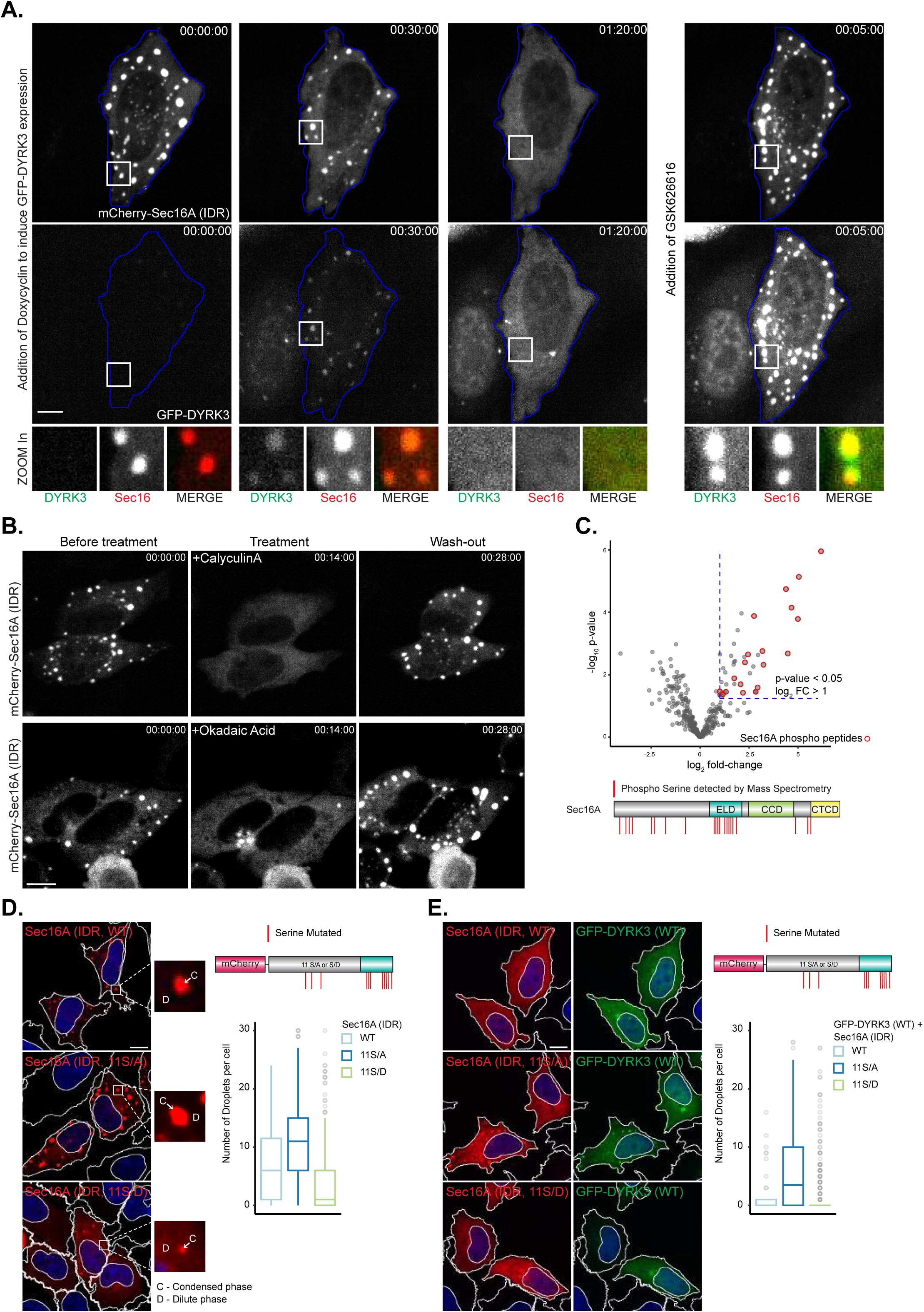
Sec16A (IDR) condensates are regulated by the opposing activity of DYRK3 kinase and Ser/Thr phosphatases. A. Time lapse images of a cell (HeLa Flp-In T-Rex) transfected with inducible GFP-DYRK3 (WT) and mCherry-Sec16A (IDR). GFP-DYRK3 expression was induced by addition of doxycycline (500 ng/mL). The dissolution of Sec16A (IDR) condensates by DYRK3 is reversed by addition of GSK626616 (1µM). B. Time lapse images of cells (HeLa Flp-In T-Rex) show dissolution of mCherry-Sec16A (IDR) upon Calyculin A (3nM) and Okadaic acid (2µM) treatment, which is reversed by washout of the drugs. C. Volcano plot shows enrichment of GFP-DYRK3 (WT) regulated phospho-sites on Sec16A, compared to GFP control. Phospho-sites with normalized log2 ratios higher than the cut-off value of 1, and p value < 0.05 (Data shown is from 4 technical replicates) are considered as DYRK3-regulated. Schematic representation of DYRK3-regulated phosphosites on full-length Sec16A is shown below. D. Left panel: Representative images of cells (HeLa) expressing mCherry-wildtype Sec16A (IDR) (Sec16A (IDR, WT)), or phospho mutants in which 11 serines were mutated to alanines (Sec16A (IDR, 11S/A)) or aspartic acid (Sec16A (IDR, 11S/D)). Condensed (C) and dilute (D) phases are marked in the inset. Right Panel (Top): Schematic representation of the serines mutated in Sec16A (IDR). Right Panel (Bottom): Quantification of condensate formation by Sec16A (IDR) mutants compared to Sec16A (IDR, WT) (Data shown is from 3 technical replicates). E. Left panel: Representative images of cells (HeLa) expressing mCherry Sec16A (IDR, WT), mCherry-Sec16A (IDR, 11S/A) or mCherry-Sec16A (IDR, 11S/D) in combination with GFP-DYRK3 (WT). Right Panel (Top): Schematic representation of the serines mutated in Sec16A (IDR). Right Panel (Bottom): Quantification of number of Sec16A (IDR, WT) or Sec16A mutants (IDR, 11S/A or 11S/D) condensates per cell, in cells expressing GFP-DYRK3 (WT) (Data shown is from 3 technical replicates). Box plots (Fig. 5D,E): centre line, median; box, interquartile range; whiskers, 1.5× interquartile range; dots, outliers. Images are representative of at least three independent experiments. All scale bars: 10 µm.

To study DYRK3-mediated phosphorylation of Sec16A we used PhosTag gels, and observed that the phosphorylation status of Sec16-IDR changes upon overexpression of DYRK3 (Suppl. Fig. 6B). A detailed analysis by phosphoproteomics revealed that at least 21phosphorylation sites were significantly enriched when full-length mCherry-Sec16A was co-expressed with GFP-DYRK3. These phosphosites were most prominently located in the N-terminal part of Sec16A (8 serines), as well as in its ER-localizing domain (ELD) (10 serines), that is part of the IDR (Fig. 5C). To test the role of these phosphosites in controlling the condensation properties of Sec16A, we generated phosphomimetic (S/D) and phosphodeficient (S/A) mutants, which revealed a synergistic effect in altering phase transition behavior. While single point mutants had no substantial effect, a Sec16A-IDR mutant in which 3 serines in the N-terminal part and 8 serines in the ELD part of the IDR were mutated into alanines (11S/A) increased the number of condensates per cell (Fig. 5D, Suppl. Fig. 6C). Consistently, the ability of DYRK3 to drive the dissolution of condensates formed by the phospho-deficient Sec16A-IDR 11S/A was significantly reduced compared to its activity on condensates generated by non-mutated Sec16-IDR (Sec16A-IDR (WT)). (Fig. 5E). Mutating the 11 serines into aspartic acid (D) increased the condensation threshold of Sec16A-IDR, as reflected in less condensates per cell and a higher concentration of the protein in the cytoplasm outside of the condensates (dilute phase). It did however not fully abolish the formation of condensates (Fig. 5D, Suppl. Fig. 6C), which could still be dissolved by DYRK3 (Fig. 5E, Suppl. Fig. 6D). This indicates that DYRK3 modulates the condensation properties of Sec16A by phosphorylating 11 serines in the Sec16A IDR and ELD, but that a full abrogation of phase separation may require the phosphorylation of additional sites in Sec16A, as well as of other components found in the Sec16A condensates that are known substrates of DYRK3, such as p115 and Sec31A(*62*).

### Sec16A phase separation controls ERES formation

Formation of protein condensates is driven by multivalent interactions, often mediated by pi-pi interactions between tyrosine-rich domains (YRDs), or by cation-pi interactions between YRDs and arginine-rich domains (RRDs) as well as by hydrophobic and CH-pi interactions of YRDs and proline-rich domains (PRDs)(*20, 63*). Since Sec16A contains both a YRD and an RRD in its ER-localizing domain (ELD) that is extensively phosphorylated by DYRK3, and some of its receptors on the ER membrane such as TANGO1L contain PRDs in their cytosolic domains(*7, 57*), its recruitment to the ER may be facilitated by phase separation. We first tested this possibility with the Sec16A-IDR condensates. Since the RRD, together with the CCD (Fig. 6A), mediates Sec16A recruitment to the ER(*11, 13, 15*), we deleted the RRD from Sec16A-IDR (mCherry-Sec16A (IDR (WT), ΔRRD). This did not abolish its ability to phase separate (Suppl. Fig. 7A). Mutating all 28 tyrosines of the YRD into glycines (mCherry-Sec16A(IDR, Y/G) and mCherry-Sec16A (IDR (Y/G), ΔRRD) however, completely prevented the formation of condensates (Fig. 6A, Suppl. Fig. 7A). Furthermore, expressing the ELD or just the YRD domain of Sec16A showed that it partitioned into Sec16A-IDR condensates, while the RRD domain by itself did not (Suppl. Fig. 7B). We next tested the PRD domain of TANGO1L, which has been shown to interact with a fragment of Sec16A containing the ELD and part of the CCD(*57*). While this domain does not form condensates by itself, it efficiently partitions into Sec16A-IDR condensates (Fig. 6B). Mutating 20 of the 45 prolines in the PRD of TANGO1L into glycines (P/G) prevented it from being recruited into Sec16A-IDR condensates, while mutating them into leucines (P/L), which can also engage in hydrophobic and CH-pi interactions, did not prevent this (Fig. 6B, Suppl. Fig. 7C) This indicates that phase separation is driven by interactions between Sec16A molecules via the YRD, and by hydrophobic and CH-pi interactions between the YRD of Sec16A and the PRD of some of its binding partners, such as TANGO1L and cTAGE5.

**Figure 6.**
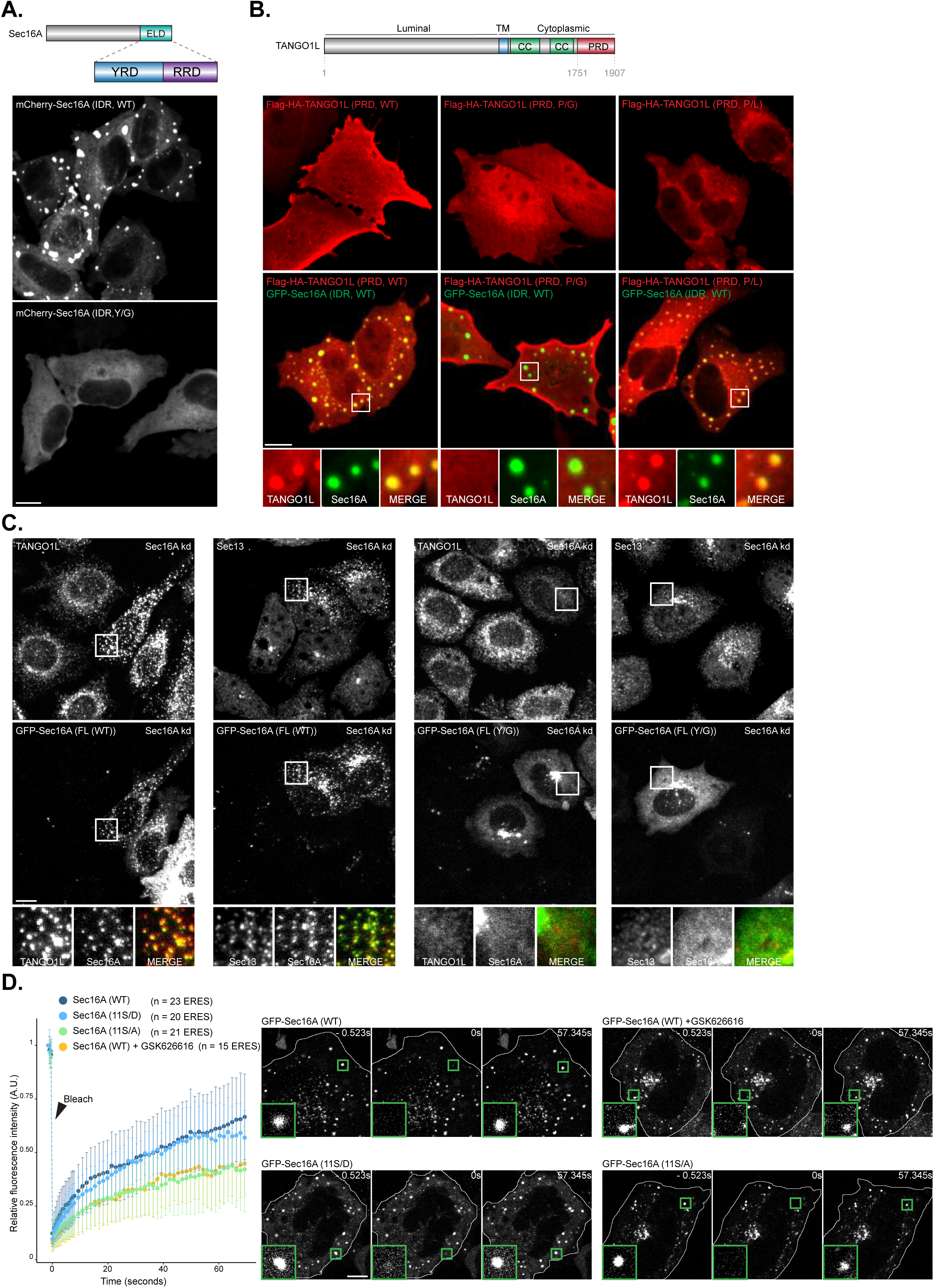
Sec16A drives ERES formation through phase separation of a tyrosine rich domain. A. Immunofluorescence images of cells (HeLa) show condensate formation by mCherry-Sec16 (IDR, WT), but not mCherry-Sec16 (IDR, Y/G). Schematic depicting domain organization of tyrosine-rich region (YRD) and arginine-rich domain (RRD) in the ER localizing domain (ELD) of Sec16A (IDR) is shown above. B. Upper Panel: Schematic depicting the domain organization of Tango1L. Lower Panel: Immunofluorescence images of cells (HeLa) show GFP-Sec16A (IDR, WT) condensates recruit Flag-HA-TANGO1L (PRD, WT) and Flag-HA-TANGO1L (PRD, P/L), but not Flag-HA-TANGO1L (PRD, P/G). C. Left Panel: Immunofluorescence images of cells (HeLa) show TANGO1L and Sec13 redistribution from ERES upon siRNA-mediated knockdown of endogenous Sec16A (Sec16A kd) is rescued by overexpression of a siRNA-resistant GFP-Sec16A (FL, WT). Right Panel: Immunofluorescence images of cells (HeLa) show TANGO1L and Sec13 redistribution from ERES upon knockdown of siRNA-mediated endogenous Sec16A (Sec16 kd) is not rescued by overexpression of a siRNA-resistant GFP-Sec16A (FL, Y/G). D. Fluorescence recovery after photobleaching (FRAP) trajectories of ERES localized GFP-Sec16A (FL, WT), GFP-Sec16A (FL, WT) in the presence of GSK626616((1µM, 1.5 h), GFP-Sec16A (FL, 11S/A) and GFP-Sec16A (FL, 11S/D) in cells (HeLa). Images show FRAP recovery. Data are mean ± s.d (The number of cells is more than 10 per condition, and 3 independent experiment. GSK626616 inhibitor data is from 2 independent experiment). Images are representative of at least three independent experiments. All scale bars: 10 µm.

To study the function of these domains in full length Sec16A, we first depleted Sec16A by RNAi (Suppl. Fig 7D), which results in a drastic change in localization of Sec13 and TANGO1L as they distribute into the reticular ER instead of accumulating at ERES(*57*) (Suppl. Fig 7E). We then asked whether RNAi-resistant forms of Sec16A could rescue the effect. While full length wild-type (WT) GFP-Sec16A rescued the formation of ERES, and co-localized with Sec13 and TANGO1L, the full length YRD (Y/G) mutant of Sec16A did not rescue ERES formation (Fig. 6C), remaining largely dissolved in the cytoplasm as well as showing some Golgi-like association. Moreover, full length Sec16A Y/G was unable to partition into TANGO1L-positive ERES in cells not depleted for endogenous Sec16A and prevented the recruitment of the cytosolic Sec13 into TANGO1L-positive ERES (Suppl. Fig. 7F). In contrast, both the 11S/A (phosphodeletion) and 11S/D (phosphomimetic) mutant of full length Sec16A still formed ERES-like structures (Suppl. Fig. 7G). To study the dynamics of their exchange between the condensed ERES and dilute cytosolic phases, we next performed photobleaching experiments. While Sec16A 11S/D displayed unaltered kinetics compared to wildtype, the 11S/A mutant displayed reduced kinetics, similar to Sec16A wildtype in the presence of GSK-626616 (Fig. 6D). Together, this indicates that Sec16A recruits ERES components such as TANGO1L to the ER by a phase separation-dependent mechanism, and that DYRK3 is critically important in regulating the dynamics of condensation and dissolution through phosphorylation of the involved components.

## Discussion

Here, we have uncovered that the dual-specificity kinase DYRK3 plays a key role in controlling the condensation of Sec16A and multiple co-condensing proteins of the early secretory pathway. Inhibiting DYRK3 activity leads to an enlargement of ERES and an accumulation of COPII budding intermediates and COPII vesicles at ERES, as well as an aberrant clustering of swollen ERGIC and cis-Golgi intermediates at these sites. Increasing the kinase activity of DYRK3 leads to a complete dissolution of ERES and to a collapse of the Golgi matrix into a spherical condensed structure. This shows that a balanced control by DYRK3 is essential for the proper functioning and turnover of these organelles and the transport of cargo between them, as well as maintaining their spatial proximity and morphology.

These findings add to the repertoire of intracellular condensates controlled by DYRK3 and extend it to secretory membrane traffic. Whether there is a cell physiological purpose for the same kinase controlling mTOR activation(*32*), mediating the dissolution of membraneless organelles during mitosis(*52*), and regulating ER export, is currently unclear. One possibility is that this reflects an actively controlled mechanism of cell size control, which depends on protein translation, surface expansion(*64, 65*), and timing of mitosis(*66, 67*). Since DYRK kinases have repeatedly been linked to cell size and cell cycle control(*66, 68–71*), and intracellular condensates have the intrinsic physical property to scale with cellular volume(*72*), this will be an interesting topic for further study. Importantly, also DYRK2 can dissolve Sec16A condensates, although less potently than DYRK3, similar to its participation in the dissolution of stress granules during stress recovery(*32*), indicating that in mammalian cells, DYRK2 and DYRK3 have partially redundant functions.

For the early secretory pathway, we have shown that DYRK3 controls LLPS through phosphorylating Sec16A, whose N-terminal half (termed Sec16A-IDR), which contains a large intrinsically disordered region (IDR) and a tyrosine-rich domain (YRD), drives LLPS. This creates a locally condensed environment in which also other ERES proteins, COPII coat proteins, and matrix proteins of the ERGIC and cis-Golgi partition, but, importantly, not medial or trans-Golgi-associated proteins. This is critically dependent on the YRD of Sec16, which localizes to the previously identified ER-localizing domain (ELD), indicating that its ER localization depends, besides the well-characterized interactions via its central conserved domain (CCD) and C-terminal conserved domain(*15*), also on the synergistic collective behavior of phase separation. The YRD achieves this by, amongst others, interacting with a proline-rich domain (PRD) within the IDR of the cytosolic domain of the co-phase-separating and ER-resident transmembrane protein TANGO1L and cTAGE5. This likely occurs by means of hydrophobic and CH-pi interactions, known to drive phase separation of other proteins(*73, 74*). Thus, through collective phase separation, driven by Sec16A and including ER-resident transmembrane proteins such as Sec12, TANGO1L and cTAGE5, locally condensed environments emerge on the cytosolic surface of the ER, from which ribosomes are excluded and in which COPII coat proteins are concentrated allowing the formation of COPII vesicles. We have also shown that matrix proteins associated with the ERGIC (TFG) and cis-Golgi (GM130 and p115) have the propensity to co-phase separate with Sec16A-IDR, yet in unperturbed cells these proteins do not fully merge. This may be due to the fact that the cis-Golgi is tightly linked to the medial-Golgi involving other golgins that do not co-phase separate with Sec16A but create their own condensate with somewhat different properties(*34, 38*). This could create a situation that is comparable to the internal organization of nucleoli and the juxtapositioning of stress granules and P-bodies, where content is exchanged without a full mixing due to different condensate properties, but with some proteins having affinity for both(*75, 76*). These opposing forces could drive a self-organized microenvironment within the cytoplasm that aligns ERES, ERGIC, and cis-Golgi without a full merging, facilitating vesicle transport. This explains the GM130 collapse into a separate condensed spherical structure when the phase-separated ERES environment is dissolved by DYRK3, as well as the well-known Brefeldin A (BFA) effect. In the latter case, the balance is disturbed in the opposite direction, where a loss of Golgi membranes due to their back-fusion with the ER allows GM130 to partition into the ERES condensate, explaining its co-localization with ERES in this condition(*77, 78*). Intriguingly, by creating an asymmetry in DYRK3 activity within the cell, as has been shown for the DYRK3 homologues Pom1 in *S. pombe* and MBK-2 in *C. elegans* (*51, 79–81*), and as suggested by DYRK3 being enriched at centrosomes(*52*), this could establish an actively controlled condensate flux, facilitating directionality of transport(*3, 41*) . It may well be that such mechanisms generally act to actively organize membrane trafficking, involving different sets of phase-separating proteins that associate with membranes, establishing local territories for the various membrane transport routes that exist in cells.

## Acknowledgements

We thank all members of the Pelkmans lab for discussions. We further thank Robin Klemm for reagents and support and critical reading of the manuscript and acknowledge the assistance and support of the Center for Microscopy and Image Analysis, University of Zurich for performing EM, FRAP, and CLEM experiments, and the Functional Genomics Center, University and ETH Zurich, for performing phosphoproteomics. EMBO and HFSP LTF supported A.K.R. L.P. is supported by the Swiss National Science Foundation and the University of Zurich.

## Author contributions

L.P. conceived the project. R.G. and A.K.R. performed all experiments and analyzed the data.

L.P., R.G. and A.K.R. interpreted the results and wrote the paper.

**Supplementary Figure 1.**
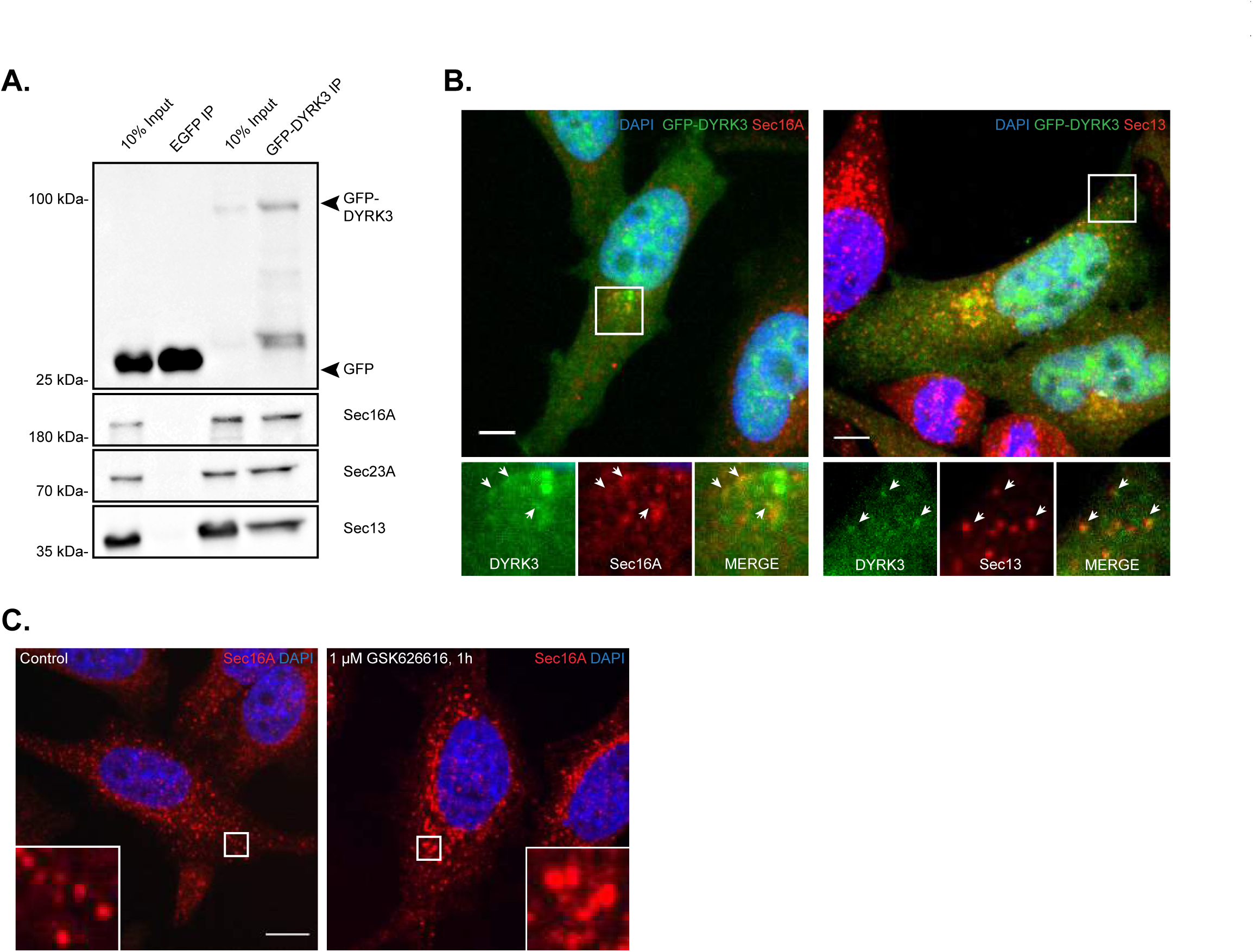
DYRK3 interacts with COPII components and regulates ER exit sites. A. Representative western blot shows GFP-DYRK3 interacts with endogenous COPII components (Sec16A, Sec23 and Sec13). HeLa cells were transfected with GFP-DYRK3 (WT) and GFP control plasmids, followed by Immuno-precipitation (IP) using GFP-Trap magnetic beads. Sec16A, Sec23A and Sec13 were probed using antibodies against the respective proteins. GFP and GFP-DYRK3 were probed with anti-GFP antibody. Lysate used for IP is marked as Input lane. B. Immunofluorescence images of cells (HeLa Flp-In T-Rex) show co-localization of GFP-DYRK3 (WT) with ERES markers (Sec16A and Sec13). Arrows indicate sites of co-localization. GFP-DYRK3 (WT) expression was induced in HeLa Flp-In T-Rex cells by adding doxycycline (500ng/mL, 4hrs) C. Immunofluorescence images of cells (HeLa) show increase in the size of ERES (Sec16A) upon GSK626616 treatment (1µM, 1h). Images are representative of at least three independent experiments. All scale bars: 10 µm

**Supplementary Figure 2.**
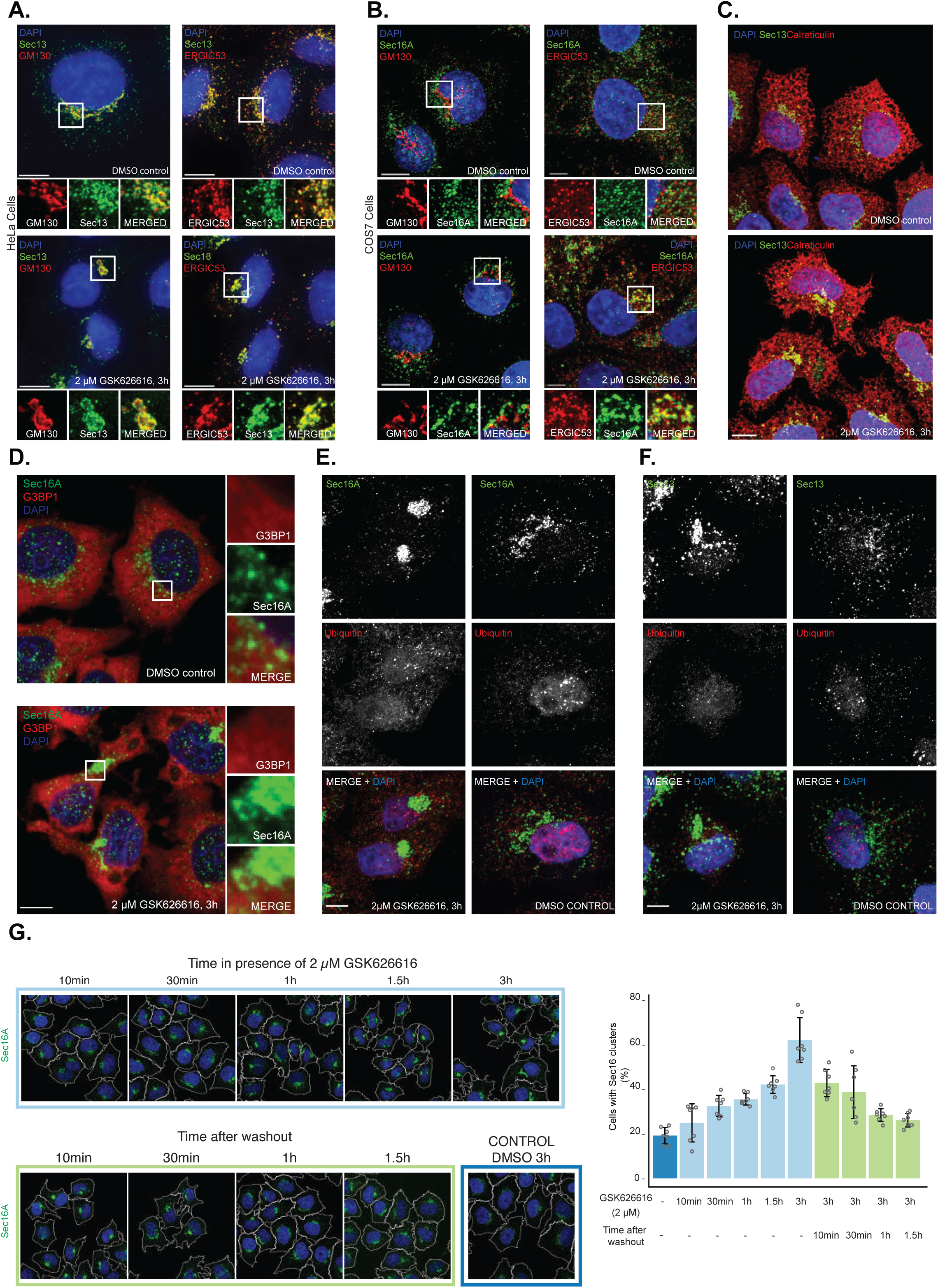
Formation of reversible perinuclear clusters of ER exit site and Golgi proteins, upon DYRK3 inhibition. A. Immunofluorescence images of cells (HeLa) stained with ERES marker (Sec13), Golgi marker (GM130) and ERGIC marker (ERGIC53) show perinuclear clustering of these components upon GSK626616 treatment (2µM, 3h). B. Immunofluorescence images of cells (COS7) stained with ERES marker (Sec16A), Golgi marker (GM130) and ERGIC marker (ERGIC53) show perinuclear clustering of these components upon GSK626616 treatment (2µM, 3h). C. Immunofluorescence images of cells (HeLa) stained with ER marker (Calreticulin) and ERES marker (Sec13) show ER network is unperturbed upon GSK626616 treatment (2µM, 3h). D. Immunofluorescence images of cells (HeLa) show the perinuclear accumulations of ERES marker (Sec16A) upon GSK626616 treatment (2µM, 3h) does not colocalize with stress granule marker (G3BP1). E. Immunofluorescence images of cells (HeLa) show the perinuclear accumulations of ERES marker (Sec16A) upon GSK626616 treatment (2µM, 3h) are not ubiquitinated aggregates (Ubiquitin). F. Immunofluorescence images of cells (HeLa) show that the perinuclear accumulations of ERES marker (Sec13) upon GSK626616 treatment (2µM, 3hr) are not ubiquitinated aggregates (Ubiquitin). G. Immunofluorescence images of cells (HeLa) show perinuclear clustering of endogenous Sec16A upon GSK626616 treatment (2µM, indicated times) is reversed upon washout of the drug. Immunofluorescence image of cells treated with DMSO for 3h is shown. Quantification of fraction of cells with perinuclear Sec16A cluster (classified by SVM training) during the time course experiment. Data are mean ± s.d (Data shown is from 3 technical replicates, 2 independent experiments). Images are representative of at least three independent experiments. All scale bars: 10 µm

**Supplementary Figure 3.**
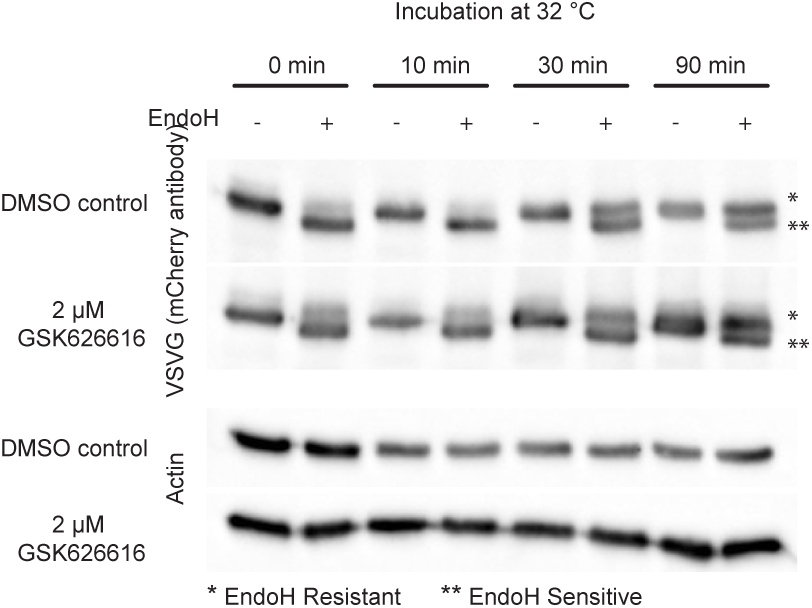
DYRK3 inhibition does not block accessibility of VSVG to Golgi enzymes. Representative western blot shows evaluation of glycosylation of VSVG (ts045-VSV-G-mCherry) by endoglycosidase H (Endo H) sensitivity assay at the permissive temperature (32°C) for indicated time points. Endo H sensitive and resistant form of VSVG is labeled in the blot. Time points represent time after shift to permissive temperature (32°C). Images are representative of at least three independent experiments.

**Supplementary Figure 4.**
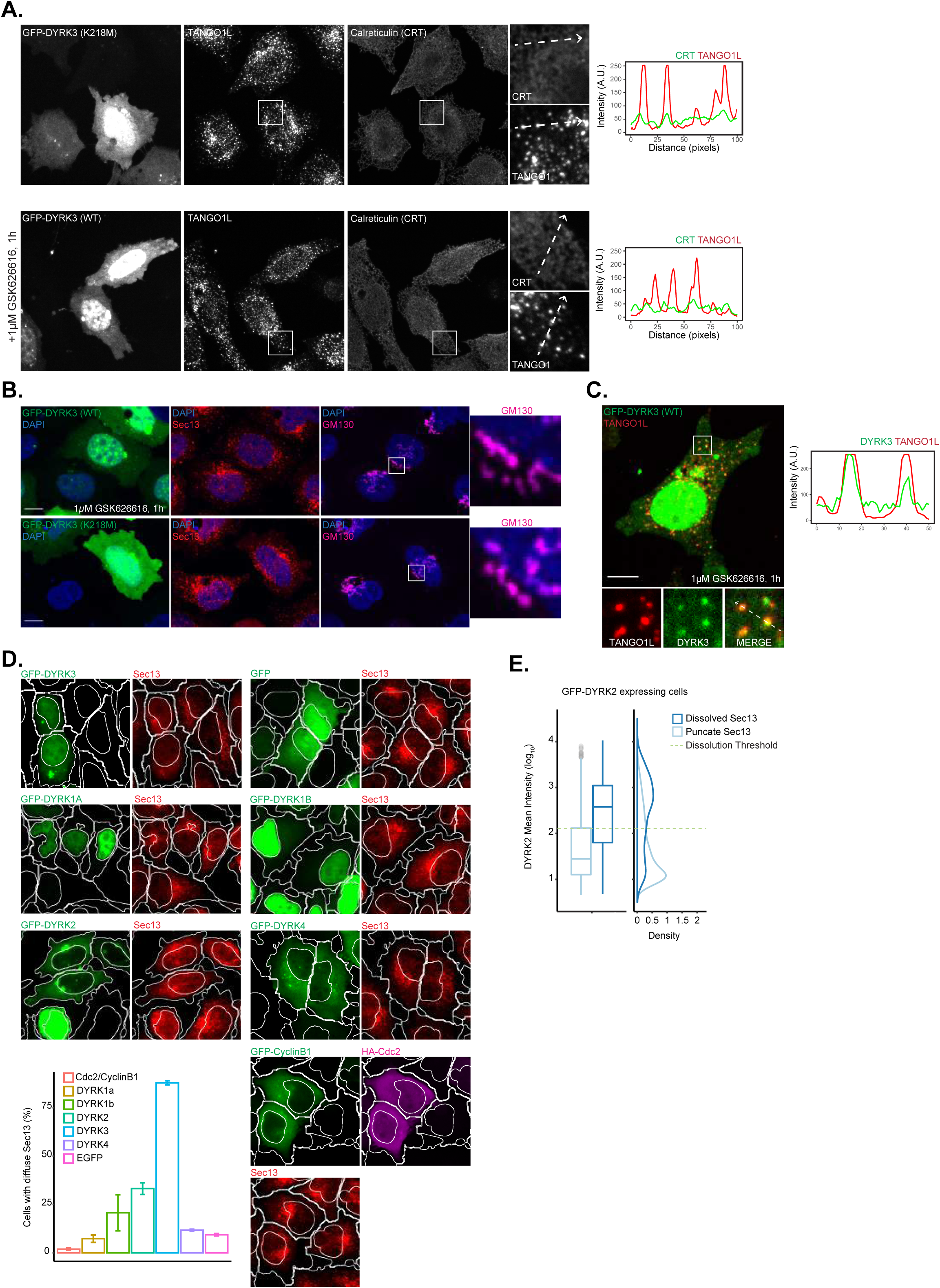
DYRK3 kinase overexpression perturbs ER exit sites and Golgi organization. A. Immunofluorescence images of cells (HeLa) show punctate TANGO1L distribution is not affected by overexpression of GFP-kinase-dead DYRK3 (GFP-DYRK3 (K218M)). The redistribution of TANGO1L across the ER network (Calreticulin (CRT)) upon overexpression of GFP-DYRK3 (WT) is reversed upon addition of GSK626616 (1µM, 1h). The fluorescent line profile of TANGO1L and CRT along the arrows in the inset is shown. B. Immunofluorescence images of cells (HeLa) show unperturbed, punctate ER exit site (Sec13) and a ribbon like Golgi structure (GM130), upon overexpression of GFP-DYRK3 (K218M). The dissolution of ER exit sites and the condensed Golgi structure observed upon overexpression of GFP-DYRK3 (WT) is reversed upon addition of GSK626616 (1µM, 1h). C. Immunofluorescence image of a cell (HeLa) shows co-localization of endogenous TANGO1L and overexpressed GFP-DYRK3 (WT) upon GSK626616 treatment (1µM, 1h). The fluorescent line profile of TANGO1L and DYRK3 along the arrows in the inset is shown. D. Immunofluorescence images of cells (HeLa) show the effect on the dissolution of Sec13 puncta (ERES) upon overexpression of DYRK family kinases and Cdc2/ CylinB1. The plot shows quantification of dissolution of Sec13 puncta by DYRK family kinases and Cdc2/CylinB1. Data are mean ± s.d (Data shown is from 3 technical replicate). E. Quantification of the dissolution of Sec13 puncta as a function of DYRK2 expression level shows concentration dependent dissolution. Cells containing a punctate or dissolved Sec13 staining in HeLa cells were classified by SVM training. Images are representative of at least three independent experiments. All scale bars: 10 µm

**Supplementary Figure 5.**
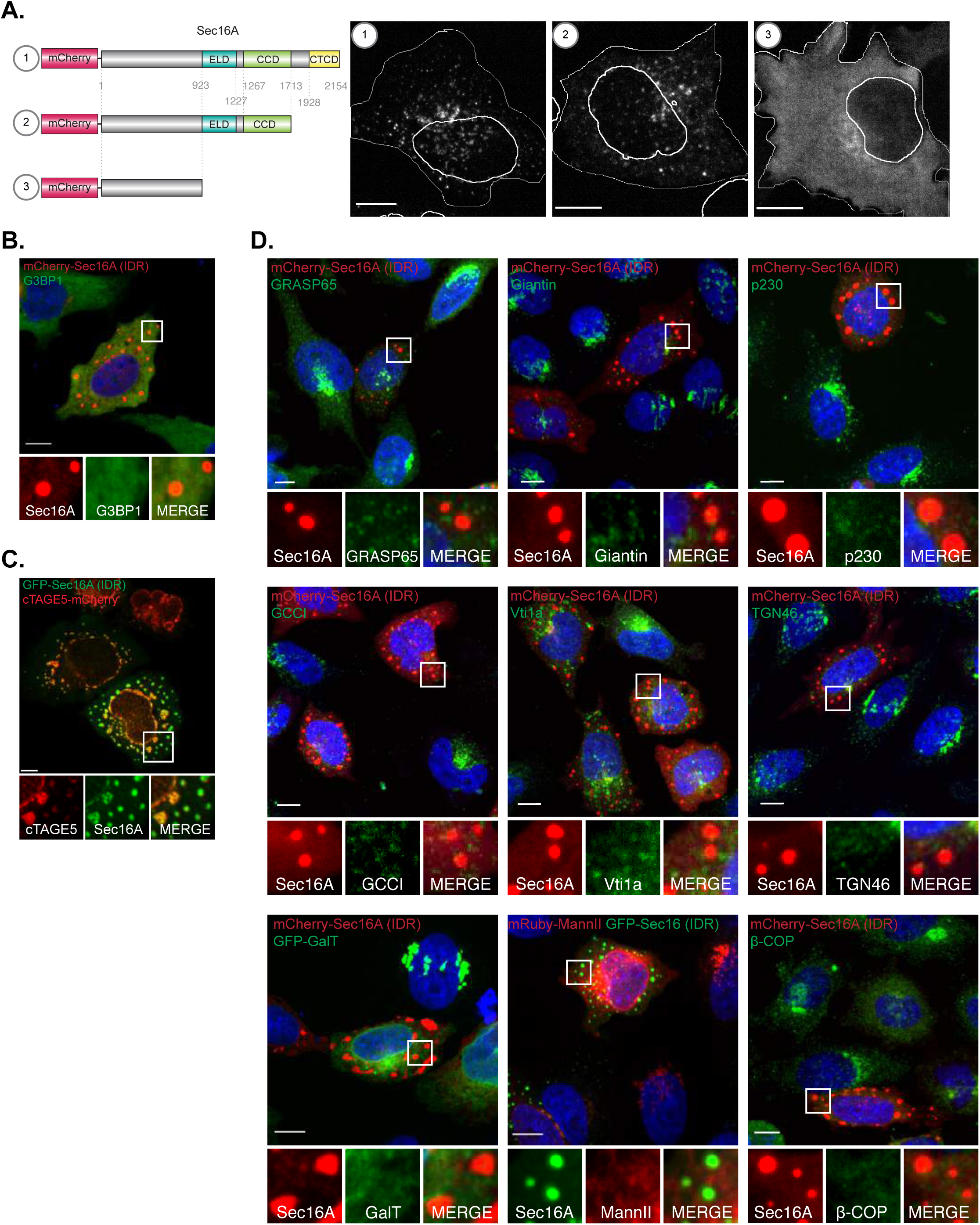
Sec16A (IDR) condensates do not recruit stress granule, medial and trans-Golgi proteins. A. Schematic representation of the mCherry-tagged full length and truncation mutants of Sec16A. Immunofluorescence images of cells (HeLa) show the localization of the full-length and truncation mutants upon overexpression. B. Immunofluorescence image of cells (HeLa) show mCherry-Sec16A (IDR) condensates do not recruit stress granule marker (G3BP1). C. Immunofluorescence image of cells (HeLa) show recruitment of overexpressed cTAGE5-mCherry to GFP-Sec16A (IDR) condensates. D. Immunofluorescence images of cells (HeLa) show mCherry- and GFP-Sec16A (IDR) condensates do not recruit Golgi stacking protein (GRASP65), medial-Golgi protein (Giantin), trans-Golgi proteins (p230, GCC1, Vti1a, TGN46), Golgi enzymes (GFP-GalT, mRuby-ManII) and COPI protein (β-COP). Images are representative of at least three independent experiments. All scale bars: 10 µm

**Supplementary Figure 6.**
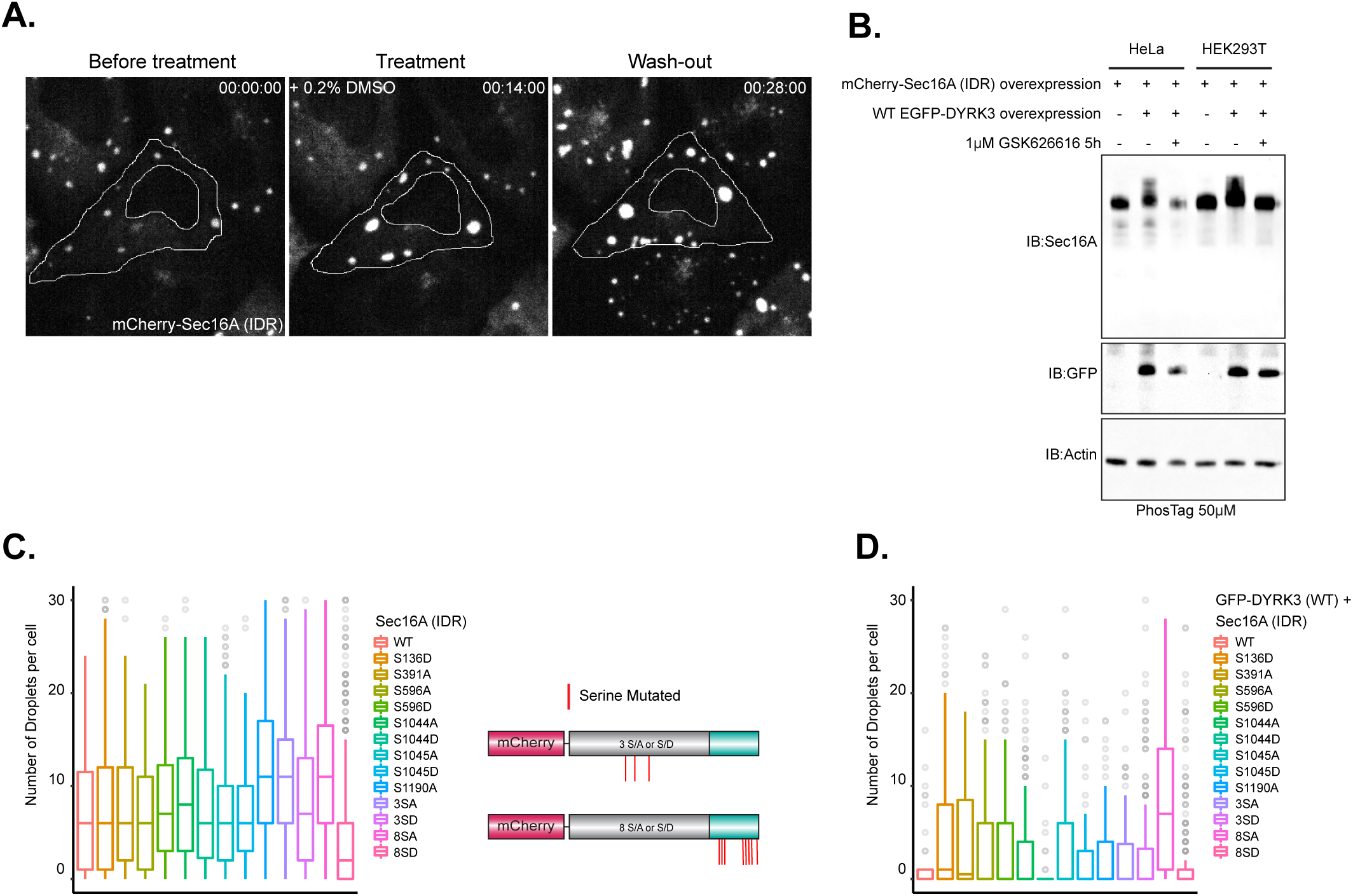
DYRK3 regulates Sec16A (IDR) condensation in a phosphorylation-dependent manner. A. Time lapse images of cells (HeLa) show mCherry-Sec16A (IDR) condensates are unaffected upon DMSO treatment. B. Representative western blot shows Phos-tag immunoblot analysis of Sec16A (IDR). Cells were transfected with mCherry-Sec16A (IDR) alone, or in combination with inducible GFP-DYRK3 (WT) in the presence or absence of GSK626616 (1µM) and run on a Phos-tag gel (50 µM). C and D. Quantification of the number of Sec16A (IDR, WT) or Sec16A mutant (IDR, S/A or S/D) condensates per cell when expressed alone (C), or in combination with GFP-DYRK3 (WT) (D). Data shown is from 3 technical replicates. Schematic depicting the phospho sites mutated in Sec16A(IDR) are shown in C. Box plots: centre line, median; box, interquartile range; whiskers, 1.5× interquartile range; dots, outliers. Images are representative of at least three independent experiments. All scale bars: 10 µm

**Supplementary Figure 7.**
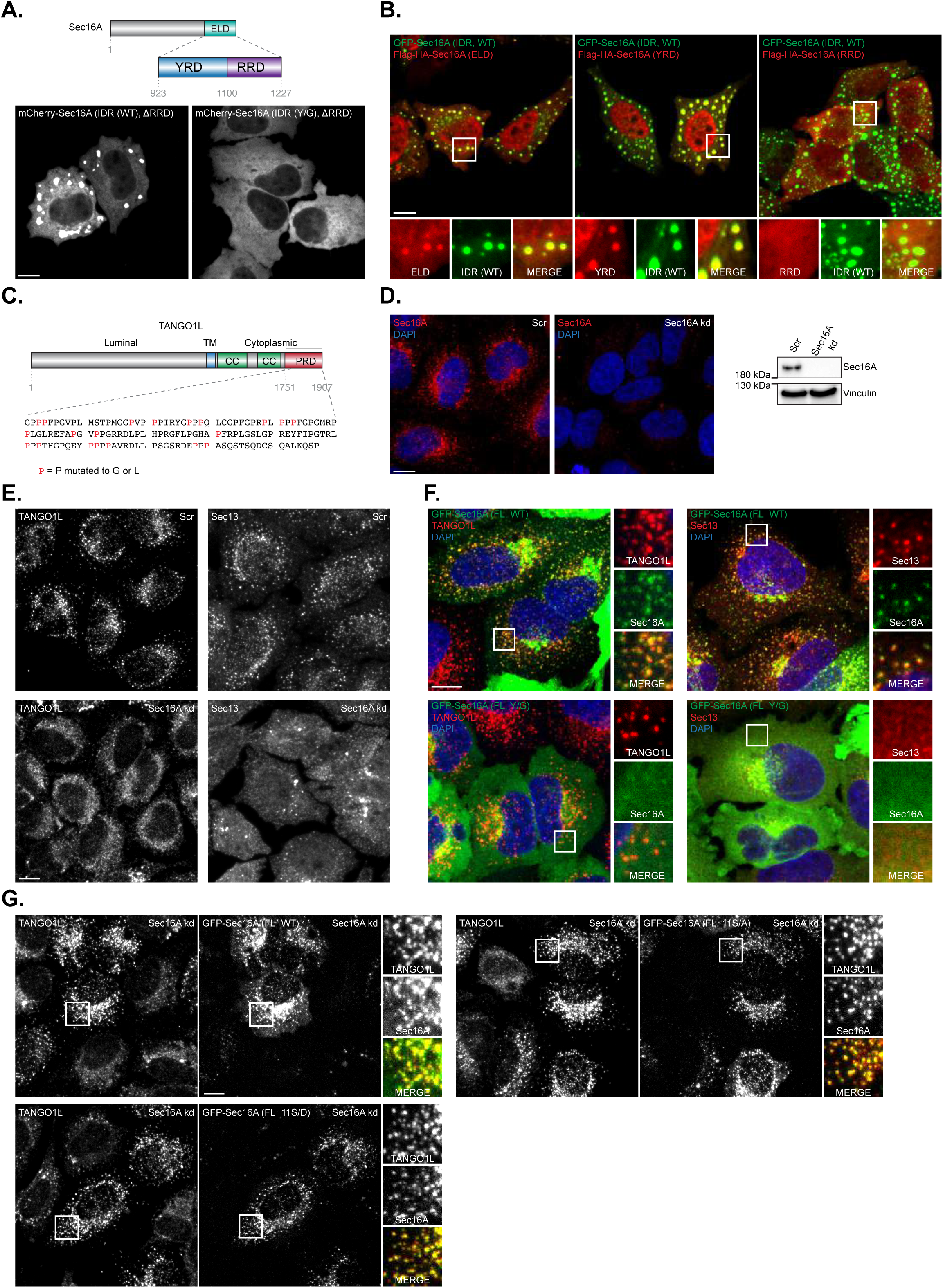
Tyrosine rich domain (YRD) in the ER localizing domain of Sec16A (IDR) is required for LLPS. A. Immunofluorescence images of cells (HeLa) show condensate formation by mCherry-Sec16 (IDR (WT), ΔRRD) but not mCherry-Sec16 (IDR (Y/G), ΔRRD). Schematic depicting domain organization of tyrosine-rich region (YRD) and arginine-rich domain (RRD) in the ER localizing domain (ELD) of Sec16A (IDR) is shown above. B. Immunofluorescence images of cells (HeLa) show GFP-Sec16 (IDR, WT) condensates recruit Flag-HA-Sec16 (ELD) and Flag-HA-Sec16 (YRD), but not Flag-HA-Sec16 (RRD). C. Schematic depicting domain organization of Tango1L and the prolines mutated in the proline rich domain (PRD). D. Immunofluorescence and western blot images show validation of siRNA-mediated knockdown of endogenous Sec16A. E. Immunofluorescence images show redistribution of Tango1L and Sec13 from ERES upon siRNA-mediated knockdown of endogenous Sec16A (Sec16 kd). F. Immunofluorescence images of cells (HeLa) show colocalization of overexpressed GFP-Sec16A (FL, WT) with ERES marker (Tango1L and Sec13). Overexpressed GFP-Sec16A (FL, Y/G) does not localize to Tango1L marked ERES, and redistributes endogenous Sec13 from ERES. G. Immunofluorescence images of cells (HeLa) show TANGO1L redistribution from ERES upon siRNA-mediated knockdown of endogenous Sec16A (Sec16 kd) is rescued by overexpression of a siRNA-resistant GFP-Sec16A (FL, WT), GFP-Sec16A (FL, 11S/A) and GFP-Sec16A (FL, 11S/D). Images are representative of at least three independent experiments. All scale bars: 10 µm

## Material and Methods

### Cell culture

HeLa cells were a kind gift from Marino Zerial (MPI-CBG, Dresden), HeLa-FlpIn-Trex cells were a kind gift from Ivan Dikic (Goethe University, Frankfurt) and HEK293T cells were from ATCC (Molsheim Cedex). Hela and HEK293T cells were maintained in 10-cm dishes in DMEM supplemented with 10% fetal bovine serum (Sigma-Aldrich) and L-Glutamine (Sigma-Aldrich). HeLa-FlpIn-Trex cells were maintained in 10-cm dishes in DMEM supplemented with 10% tetracycline-free fetal bovine serum (Clonetech Laboratories), L-Glutamine and blasticidin (1µg/mL, Santa Cruz). Hela-FlpIn-Trex expressing GFP-wildtype DYRK3 was induced using doxycycline (Sigma) with indicated time and concentration. All cells were maintained in a humidified incubator at 37°C under 5% CO_2_. For imaging experiments, cells were grown in 96-well plates (Greiner Bio-One International).

### Immunofluorescence

Cells were grown in 96-well plates and fixed by adding 4% PFA (Electron Microscopy Sciences) for 20 mins and then permeabilized with 0.25% Titon-X-100 for 10 mins. The cells were blocked with 1% BSA in PBS (blocking buffer) before incubating with primary antibodies in blocking buffer for 2 hrs at room temperature, followed by incubation with Alexa-Fluor labeled secondary antibody (Life Technologies) in PBS for 1 hour at room temperature. Nuclei were stained using DAPI (Life Technologies).

### Antibodies

The following antibodies were used: Sec16A (1:300, Bethyl Laboratories, A300-648A), Sec16A (1:400, Thermo Fisher Scientific, PA5-52182), Sec13 (1:1000, Bethyl Laboratories, A303-980A), Sec23A (1:300, Thermo Fisher Scientific, PA1-069A), Actin (1:1000, Abcam, ab3280), GFP (1:1000, Cell Signaling Technologies, 2956S), GM130 (1:500, BD Biosciences, 610823), ERGIC53 (1:400, Enzo Life Sciences, ABS300-0100), G3BP1 (1:400, Abcam, ab56574), Ubiquitin (FK2) (1:200, Enzo Life Sciences, BML-PW8810), mCherry (1:1000, Novus Biologicals, NBP2-25157), MIA3 (1:400, Sigma-Aldrich, HPA055922), HA (1:500, Abcam, ab130275), TFG (1:200, Sigma-Aldrich, HPA019473), p115 (1:300, Thermi Fisher Scientific, PA1006), Sec12 (1:200, Novus Biologicals, NBP1-87056), GRASP65 (1:500, Thermo Fisher Scientific, PA3910), Giantin (1:1000, Abcam, ab174655), p230 (1:300, BD Biosciences, 611280), GCC1 (1:500, Sigma-Aldrich, HPA021323), Vti1a (1:200, BD Biosciences, 611220), TGN46 (1:600, Sigma-Aldrich, T7576), β-COP (1:300, Novus Biologicals, NBP2-22544), Flag (1:800, Sigma-Aldrich, F1804), Vinculin (1:1000, Sigma-Aldrich, V9131), Calreticulin (1:200, kindly provided by Robin Klemm, UZH, Zurich, Switzerland).

### Plasmids Transfection

Plasmids transfections were performed with indicated plasmids using Lipofectamine 2000 (ThermoFisher) or genejuice reagent (EMD Millipore) as per manufacturer instructions.

### Plasmids

Generation of full-length DYRK3 plasmids, pcDNA5-GFP-wildtype DYRK3 (GFP-DYRK3 (WT)) and pcDNA5-GFP-K218M DYRK3 (GFP-DYRK3 (K218M)) has been reported earlier(*32*). Generation of pcDNA5/FRT/TO-GFP-wildtype DYRK3 (inducible GFP-DYRK3 (WT)) vector has been reported earlier(*52*). Full-length GFP-Sec16A was a kind gift from Benjamin Glick (Addgene plasmid # 15776)(*58*). Full-length and truncation mutants of Sec16A were cloned into mCherry2-C1 plasmid (kind gift from Michael Davidson, Addgene plasmid # 54563) using EcoRI and KpnI restriction sites. Primer pairs for amplifying mCherry-full-length Sec16A: 5’-TTCGAATTCTA TGCCTGG GCTCGACCG-3’ and 5’-CGCGGTACCTTATTAGTTCAGC ACCAGGTGCTTC CTC-3’; for mCherry-Sec16A (1-1227) named mCherry-Sec16A (IDR): 5’-TTCGAATTCTAT GCCTGGGCTCGA CCG-3’ and 5’-CGCGGTACCTTATTAG TAGGCAAAATCGCCGTGAA AGGAG-3’; for mCherry-Sec16A (1-1713): 5’-TTCGAA TTCTATGCCTGGGCTCGACCG-3’ and 5’-CGCGGTACCTTATTAAGCCCCCTCCTTAATCTGCCGCTC-3’; for mCherry-Sec16A (1-923): 5’-TTCGAATTCTATGCCTGGGCTCGACCG-3’ and 5’-CGCGGTACC TTATTATGCTGGTGACTGT GCTGAGTTC -3’. GFP-Sec16A (IDR) was generated by amplifying Sec16A (1-1227) from full-length GFP-Sec16A using primer pair 5’-T T C G A A T T C T A T G C C T G G G C T C G A C C G - 3’ a n d 5’ - CGCGGTACCTTATTAGTAGGCAAAATCGCCGTGAAAGGAG-3’, followed by insertion into mEGFP-C1 plasmid (kind gift from Michael Davidson, Addgene plasmid # 54759) using EcoRI and KpnI restriction sites. pEGFP-VSVG was a gift from Jennifer Lippincott-Schwartz (Addgene plasmid # 11912)(*56*). mCherry-VSVG was generated by amplifying V S V G f r o m p E G F P - V S V G u s i n g p r i m e r p a i r 5’ - TA G C T C G A G C G C C A C C A T G A A G T G C C T T T T G - 3’ a n d 5’ - GACGAATTCGTCCAAGTCGGTT CATCTCTATGTCTG-3’, followed by insertion into mCherry2-N1 plasmid (kind gift from Michael Davidson, Addgene plasmid # 54517) using EcoRI and XhoI restriction sites. EGFP-GalT was a gift from Jennifer Lippincott-Schwartz (Addgene plasmid # 11929)(*82*).

mRuby2-MannII-N-10 was a gift from Michael Davidson (Addgene plasmid # 55903)(*83*). Sec16A fragment with eight serines in the ELD mutated to alanines (S/A) or aspartates (S/D) was synthesized by Genscript (USA) and inserted into mCherry-Sec16A (IDR) using restriction sites AflII and KpnI. Additional S/A and S/D mutations were generated by using Q5^®^ Site-Directed Mutagenesis Kit (NEB, E0554S) following manufacturer’s instructions.

Sec16A fragment with twenty-eight tyrosines in the Y-rich domain of ELD mutated to glycines was synthesized by Genscript (USA) and inserted into mCherry-Sec16A (1-1227, IDR) using AflII and SanDI restriction sites, to generate mCherry-Sec16A (IDR, Y/G). mCherry-Sec16A (IDR (WT), ΔRRD) was generated by amplifying Sec16A fragment from full-length GFP-Sec16A using primer pair 5’-GACGGTACCATGCCTGGGCTCGACCG-3’ and 5’-GGTGGATCCACTGTAGTGGTAACGATCCCAGTGAC-3’, followed by insertion into mCherry2-C1 plasmid using KpnI and BamHI restriction sites. mCherry-Sec16A (IDR (Y/G), ΔRRD) mutant was generated by amplifying Sec16A fragment from mCherry-Sec16A (IDR, Y/G) using primer pair 5’-TTCGAATTCTATGCCTGGGCTCGACCG-3’ and 5’-CGCGGTACCTTATTAACTGCCGTGGCCACGATC-3’, followed by insertion into mCherry2-C1 plasmid using EcoRI and KpnI restriction sites. ER localizing domain (ELD), tyrosine rich domain (YRD) and arginine rich domain (RRD) were amplified from full-length G F P - S e c 1 6 A u s i n g p r i m e r p a i r s 5’ - C T G G A A T T C T G A G T C T G G T T C T G G T C G A C G C G - 3’ a n d 5’ - ATGCTCGAGTTATTAGTAGGCAAAATCGCCGTGAAAGGAG-3’; primer pairs 5’- C T G G A A T T C T G A G T C T G G T T C T G G T C G A C G C G - 3’ a n d 5’ - ATGCTCGAGTTATTAACTGTAGTGGTAACGATCCCAGTG-3’; primer pairs5’-C T G G A A T T C T G G C T A G A G T C A G G G A C C C C C G - 3’ a n d 5’ - ATGCTCGAGTTATTAGTAGGCAAAATCGCCGTGAAAGGAG-3’, respectively. The amplicons were inserted into pcDNA3 Flag HA plasmid (kind gift from William Sellers, Addgene plasmid # 10792) using EcoRI and XhoI restriction sites, to generate Flag-HA-Sec16A (ELD), Flag-HA-Sec16A (YRD) and Flag-HA-Sec16A (RRD).

Tango1L (PRD) wildtype (WT) fragment, and the fragments with mutation of twenty prolines into glycines (P/G) or leucines (P/L) were synthesized by Genscript (USA) and inserted into pcDNA3 Flag-HA plasmid using BamHI and EcoRI restriction sites, to generate Flag-HA-TANGO1L (PRD, WT), Flag-HA-TANGO1L (PRD, P/G) and Flag-HA-TANGO1L (PRD, P/ L).

Full length GFP-Sec16A with S/A and S/D mutations were generated by assembling N-terminal 11S/A and 11S/D Sec16A fragment from mCherry-Sec16A (1-1227, IDR) mutants, and C-terminal Sec16A (1228-2154) fragment from full-length wild type GFP-Sec16A, using Gibson Assembly^®^ Cloning Kit (NEB, E5510S) following manufacturer’s instructions. RNAi-resistant forms of wildtype, phosphospho-deletion (11S/A) and phospho-mimetic mutants (11S/D) of full-length SEC16A were generated by mutating 10 nucleotides in the silencer-select Sec16A siRNA (5’-CGGAGCUUCUGUUACGAGATT-3’, siRNA id: s19236, Thermo-Fischer Scientific) binding site using Q5^®^ Site-Directed Mutagenesis Kit (NEB, E0554S) following manufacturer’s instructions.

### Imaging

Imaging was carried out on an automated spinning disk microscope from Yokogawa (Cell Voyager 7000), with an enhanced CSU-X1 spinning disk (Microlens-enhanced dual Nipkow disk confocal scanner, wide view type), a 40× 0.95 NA Olympus objective, a 60x 1.2 NA Olympus water-immersion objective and Andor sCMOS cameras (Andor, 2,560 × 2,160 pixels). Live cell imaging experiments were performed with Yokogawa (Cell Voyager 7000) with 60x 1.2 NA Olympus water-immersion objective or with a spinning disk microscope from Nikon (Eclipse Ti-E) with an enhanced CSU-W1 spinning disk (Microlens-enhanced dual Nipkow disk confocal scanner, wide view type), a Nikon CFI PlanApo 100x oil immersion objective of NA 1.49, and a Photometric Prime BSI sCMOS (2,048 × 2,048 pixels). All cells were maintained in a humidified environment at 37^0^C under 5% CO2 for live imaging experiments.

### Western Blotting

The protein concentration in in cell lysates were determined with the Protein-660 assay containing when necessary the ionic detergent compatibility reagent (Pierce, Thermo-Fisher). The samples were denatured by the addition of SDS sample buffer (62.5 mM Tris-HCl pH 6.8; 25% glycerol; 2% SDS. 0.5% bromophenol blue) and heated at 95 °C for 10 min. The proteins were separated using a 8-12% SDS-PAGE and analysed by immunoblotting. The proteins were transferred onto a polyvinylidene fluoride (PVDF) membrane by semi-dry transfer using the Trans-Blot turbo RTA Mini PVDF transfer kit (Bio-Rad) and the Trans-Blot turbo system. After the transfer the membranes were incubated in 5% Milk in PBS for 1 h at room temperature. The membranes were then incubated with primary antibodies specific for the proteins of interest at 1:1000 dilution in 5% Milk in PBS over night at 4 °C or at room temperature for 2 hours. The membranes were washed 3 x with PBS containing 1% Tween-20 and incubated for 60 min with secondary antibodies coupled to horse-radish peroxidase. The proteins of interest were then developed by enhanced chemiluminescence (ECL, GE) and visualized using an ECL detection system (Vilber Lourmat). The intensities of the bands were quantified using ImageJ.

### Ts045-VSV-G traffic monitoring

HeLa cells were seeded onto Greiner 96-well plates, grown to ∼70% confluence and transiently transfected with the mutated ts045-VSV-G fused to GFP- or mCherry-tag using Lipofectamine 2000. The cells were kept at 37°C for 1h, then incubated at non-permissive temperature (40°C) overnight. Ts045-VSV-G trafficking was initiated by shifting to the permissive temperature of 32°C for different length of time, as specified in the figures, before fixation. When necessary, reagents were diluted in pre-warmed media and then added to the cells an hour prior switch to 32°C.

### Endoglycosidase H digestion

HeLa cells were grown in 6-cm plates. When 70 % confluence was achieve, the cells were transfected with different plasmids or plasmid combinations as reported in the figures, kept at 37°C for 1h, then incubated at 40°C. The following day, the cells were incubated at 32°C for the indicated times before lysing in RIPA lysis buffer. The samples were clarified by centrifugation at 10,000 g for 12 min and Endoglycosidase H (Endo H; New England Biolabs, Ipswich, MA, USA) digestion was performed with each sample, according to the instruction provided by the manufacturer and previously reported(*84*). Briefly, 15 µg of total protein lysate, in a 9 µl reaction volume, were mixed with 1 ml of denaturing buffer and boiled for 10 min at 100°C. After that, 2 µl of Endo H buffer, 0.5 µl of Endo H, and 7.5 µl of H2O were added to the denatured reaction and incubated at 37°C for 18 h. Western blot analysis was performed as previously above.

### Phos-tag experiments

Cells were transfected with indicated plasmids for 24 hr. Cells were washed thrice with wash buffer (10mM Tris-HCl, 150mM NaCl, pH=7.4) and lysed in Lysis buffer (50mM Tris, 150mM NaCl, 1mM DTT, 0.5% Triton-X, 2x Protease inhibitor and 2x Phosphatase inhibitor, pH=7.4) by passing five times through an insulin syringe. Lysates were spun at 20,000 g for 10 min and supernatants were stored at -80^0^C. Protein samples were run on Phos-tag gels (7% SDS-PAGE with 50 µM Phos-tag and 100 µM Zn(NO_3_)_2_, Phos-tag (TM) Acrylamide AAL-107, Wako Chemicals) in running buffer (50 mM MOPS, 50 mM Tris Base, 0.1% SDS, pH 7.8, 5 mM sodium bisulfite). Gels were then washed twice (30 min each) in transfer buffer containing 1 mM EDTA (1×NuPAGE Transfer Buffer, Invitrogen, NP00061). Western blot transfer on PVDF membranes was performed overnight at 4°C in 1×NuPAGE Transfer Buffer containing 10% methanol and 5 mM sodium bisulfite, at 4°C. PVDF membranes were further processed as described in the Western blotting section.

### Immunoprecipitation experiments

Cells were collected in lysis buffer (150 mM NaCl, 50 mM HEPES, 1% Triton X-100, 0.1% SDS, 5mM EDTA and 2x protease inhibitor cocktail in miliQ H2O), allowed to lyse on ice for 20 min. The cells were then centrifuged at 21,000 RCF for 15 min at 4^0^C and the supernatant was used for Immunoprecipitation experiments. The cleared lysates was diluted 10 fold in dilution buffer (150 mM NaCl, 50 mM HEPES and 5mM EDTA and 1x protease inhibitor cocktail), and then incubated with anti-GFP magnetic agarose beads (Chromotek) at 4 °C for 90 min on a rotory shaker. The beads were then washed three times with wash buffer (150 mM NaCl, 50 mM HEPES and 5mM EDTA). To pull down the immunocomplexes, beads were boiled in 30 µl of 2× SDS-PAGE sample buffer for 15 mins. The immunoprecipitated proteins were separated by SDS-PAGE. Western blot analysis was performed as previously described.

### Fluorescence recovery after photobleaching (FRAP) experiments

HeLa-FlpIn-Trex cells were grown on 8-well chambers (Ibidi GmbH, Germany).

Cells were transfected with the indicated plasmids for 12-16 hr before FRAP experiments. For experiments with mCherry-Sec16A(IDR), cells were imaged in DMEM supplemented with 10% fetal bovine serum (Sigma-Aldrich). For experiments with GFP-Sec16A (FL, WT), Sec16A (FL, WT) in the presence of GSK626616, GFP-Sec16A (FL, 11S/A) and GFP-Sec16A (FL, 11S/D), cells were washed thrice with DMEM without fetal bovine serum and imaged in the same medium an hour later. GSK626616 treatment (1µM) was performed in DMEM without fetal bovine serum for 1.5 hr before imaging. Photobleaching was performed on a Leica SP5 Mid UV-VIS equipped with a 63× 1.4 NA, oil, Plan-Apochromat objective. Cells were maintained at 37 °C under 5% CO_2_ during the course of experiment. Cells were imaged for 1.5 seconds, followed by photobleaching a defined region that encircles the intracellular condensate or ERES thrice at full laser power, and then recovery was monitored for approximately 40-70 seconds. Recovery curves are plotted for condensates and ERES that remained in focus (single imaging plane) during the time lapse acquisition. FRAP analysis was performed using Fiji software(*85*).

### siRNAs Transfection

For siRNAs transfection about 2800 cells were plated per well in 96-well plates (Greiner) for reverse transfection on top of a mixture of siRNAs (5nM final concentration) and RNAiMAX (0.04 µl per well in OptiMEM; ThermoFisher) according to manufacturer’s specifications. Cells were subsequently grown for 72 hours at 37°C in complete DMEM to establish efficient knock down of the targeted genes. Silencer-select siRNA for Sec16A (5’-CGGAGCUUCUGUUACGAGATT-3’, siRNA id: s19236), positive control silencer-select siRNA against KIF11 and negative control silencer-select siRNA were from Thermo-Fischer Scientific.

### Phosphoproteomics

HeLa cells were grown in 10-cm dishes to 70% confluence, transfected with 4 µg of mCherry-Sec16A plasmid in combination with 4 µg of GFP-DYRK3 (WT) or EGFP-C1 plasmids using Lipofectamine 2000 and 18h post-transfections cells were processed for immunoprecipitation as described above, using RFP-trap magnetic beads (Chromotek). The samples were then processed for on-beads digestion and phosphopeptide enrichment. For each sample, the washed beads were re-suspended in 45 µl digestion buffer (10 mM Tris + 2 mM CaCl2, pH 8.2) and the proteins were on-bead digested using 5 µl of Sequencing Grade Trypsin (100 ng/µl in 10 mM HCl, Promega). The digestion was carried out at 37°C over night. The supernatants were transferred in new tubes and the beads were washed with 150 µl TFA-buffer (0.1% TFA, 50% acetonitrile). These supernatants were collected and combined with the previously collected one. The samples were dried almost to completeness (∼5 µl) for enrichment of the phosphopeptides.

The phosphopeptide enrichment was performed using a KingFisher Flex System (Thermo Fisher Scientific) and MagReSyn Ti-IMAC beads (ReSyn Biosciences, Gauteng, South Africa)(*86*). Beads were conditioned following the manufacturer’s instructions, consisting of 2 washes with 200 µl of 70% ethanol, 1 wash with 100 µl of 1 M NH_4_OH and 3 washes with loading buffer (0.1 M glycolic acid in 80% ACN, 5% trilfuoroacetic acid (TFA)). Samples were diluted with 200 µl of loading buffer. The beads, wash solutions and the samples were loaded into 96 deep well plates and transferred to the KingFisher. The protocol of the robot carried out the following steps: washing of the magnetic beads in loading buffer (5 min), binding of the phosphopeptides to the beads (20 min), washing the beads in wash1 (0.1 M glycolic acid in 80% ACN, 5% TFA, 2 min), whash2 (80% ACN, 1% TFA, 2 min), wash3 (10% ACN, 0.2% TFA, 2 min) and eluting the phosphopeptides from the magnetic beads (1 M NH_4_OH, 10 min). The phosphopeptides were dried to completeness and re-solubilized with 10 µl of 3% acetonitrile, 0.1% formic acid for MS analysis.

### Liquid chromatography-mass spectrometry analysis

The mass spectrometry analysis of the pull-down samples was performed on a Q Exactive HF-X mass spectrometer (Thermo Scientific) equipped with a Digital PicoView source (New Objective) and coupled to a M-Class UPLC (Waters). Solvent composition at the two channels was 0.1% formic acid for channel A and 0.1% formic acid, 99.9% acetonitrile for channel B. For each sample 2 µl of peptides were loaded on a commercial MZ Symmetry C18 Trap Column (100Å, 5 µm, 180 µm x 20 mm, Waters) followed by nanoEase MZ C18 HSS T3 Column (100Å, 1.8 µm, 75 µm x 250 mm, Waters). The peptides were eluted at a flow rate of 300 nL/min by a gradient from 8 to 27% B in 85 min, 35% B in 5 min and 80% B in 1 min. Samples were acquired in a randomized order. The mass spectrometer was operated in data-dependent mode (DDA), acquiring a full-scan MS spectra (350−1’400 m/z) at a resolution of 120’000 at 200 m/z after accumulation to a target value of 3’000’000, followed by HCD (higher-energy collision dissociation) fragmentation on the twenty most intense signals per cycle. HCD spectra were acquired at a resolution of 15’000 using a normalized collision energy of 25 and a maximum injection time of 22 ms. The automatic gain control (AGC) was set to 100’000 ions. Charge state screening was enabled. Singly, unassigned, and charge states higher than seven were rejected. Only precursors with intensity above 250’000 were selected for MS/MS. Precursor masses previously selected for MS/MS measurement were excluded from further selection for 30 s, and the exclusion window was set at 10 ppm. The samples were acquired using internal lock mass calibration on m/z 371.1012 and 445.1200.

The mass spectrometry proteomics data were handled using the local laboratory information management system (LIMS)(*87*).

The mass spectrometry analysis of the phosphoproteomics samples was also performed on a Q Exactive HF-X mass spectrometer (Thermo Scientific) equipped with a Digital PicoView source (New Objective) and coupled to a M-Class UPLC (Waters). Solvent composition at the two channels was 0.1% formic acid for channel A and 0.1% formic acid, 99.9% acetonitrile for channel B. For each sample 4 µl of peptides were loaded on a commercial MZ Symmetry C18 Trap Column (100Å, 5 µm, 180 µm x 20 mm, Waters) followed by nanoEase MZ C18 HSS T3 Column (100Å, 1.8 µm, 75 µm x 250 mm, Waters). The peptides were eluted at a flow rate of 300 nL/min by a gradient from 8 to 27% B in 55 min, 35% B in 5 min and 80% B in 1 min. Samples were acquired in a randomized order. The mass spectrometer was operated in data-dependent mode (DDA), acquiring a full-scan MS spectra (350−1’400 m/z) at a resolution of 120’000 at 200 m/z after accumulation to a target value of 3’000’000, followed by HCD (higher-energy collision dissociation) fragmentation on the ten most intense signals per cycle. HCD spectra were acquired at a resolution of 60’000 using a normalized collision energy of 25 and a maximum injection time of 120 ms. The automatic gain control (AGC) was set to 1’000’000 ions. Charge state screening was enabled. Singly, unassigned, and charge states higher than seven were rejected. Only precursors with intensity above 100’000 were selected for MS/MS. Precursor masses previously selected for MS/MS measurement were excluded from further selection for 30 s, and the exclusion window was set at 10 ppm. The samples were acquired using internal lock mass calibration on m/z 371.1012 and 445.1200.

The mass spectrometry proteomics data were handled using the local laboratory information management system (LIMS)(*87*).

### Protein and phosphopeptide identification, quantification and integrative analysis

Individual data analysis workflows have been used for global protein and phosphopeptide analysis. The statistical analysis was done in a two-group comparison such as: Dyrk3 vs GFP. The acquired raw MS data were processed by MaxQuant (version 1.6.2.3), followed by protein identification using the integrated Andromeda search engine(*88*). Spectra were searched against a Uniprot human reference proteome (taxonomy 9606, canonical version from 2016-12-09), concatenated to its reversed decoyed fasta database and common protein contaminants. Carbamidomethylation of cysteine was set as fixed modification, while methionine oxidation and N-terminal protein acetylation were set as variable. Enzyme specificity was set to trypsin/P allowing a minimal peptide length of 7 amino acids and a maximum of two missed-cleavages. MaxQuant Orbitrap default search settings were used. The maximum false discovery rate (FDR) was set to 0.01 for peptides and 0.05 for proteins. Label free quantification was enabled and a 2 minutes window for match between runs was applied. In the MaxQuant experimental design, each file is kept separate in the experimental design to obtain individual quantitative values. Protein fold changes were computed based on Intensity values reported in the proteinGroups.txt file. A set of functions implemented in the R package SRMService (http://github.com/protViz/SRMService) was used to filter for proteins with 2 or more peptides allowing for a maximum of 4 missing values, normalizing the data with a modified robust z-score transformation and computing p-values using the moderated t-test with pooled variance (as implemented in the limma package(*89*)). If all measurements of a protein are missing in one of the conditions, a pseudo fold change was computed replacing the missing group average by the mean of 10% smallest protein intensities in that condition. The acquired raw MS data were processed by MaxQuant (version 1.6.2.3), followed by protein identification using the integrated Andromeda search engine(*88*). Each file is kept separate in the experimental design to obtain individual quantitative values. Spectra were searched against the same database used for proteome analysis. Carbamidomethylation of cysteine was set as fixed modification, while serine, threonine and tyrosine phosphorylation, methionine oxidation and N-terminal protein acetylation were set as variable. Enzyme specificity was set to trypsin/P allowing a minimal peptide length of 7 amino acids and a maximum of two missed-cleavages. Precursor and fragment tolerance was set to 10 ppm and 20 ppm, respectively for the initial search. The maximum false discovery rate (FDR) was set to 0.01 for peptides and 0.05 for proteins. Label free quantification was enabled and a 2 minutes window for match between runs was applied. The re-quantify option was selected. For the phospho-site analysis a similar data analysis strategy as described in (*90*) was implemented. In brief, the MaxQuant phospho_STY_site.txt file was used as the input file. The phospho-site table is expanded with respect to their multiplicity and filtered for a minimum localization site probability of 0.75. For each two-group comparison all peptides with a maximum of four missing values were retained. The data (like for the total proteome) was normalized with a modified robust z-score transformation and p-values were computed with a moderated t-test with pooled variance (as implemented in the limma package(*89*)). If all measurements of a protein are missing in one of the conditions, a pseudo fold change was computed replacing the missing group average by the mean of 10% smallest protein intensities in that condition. Calculated p-values are adjusted for multiple testing using BH-method. The statistics of the phospho peptide analysis and the total proteome analysis were merged together.

### Electron microscopy-chemical fixation

HeLa cells were grown onto 6 mm sapphire discs that were coated with L-polylysine (100 µm thickness, Engineering Office M. Wohlwend, Sennwald, Switzerland, a kind gift by Drs. Urs Ziegler and Andres Käch at the center of microscopy of the University of Zurich, ZMB-UZH), and were gold sputter labeled with fiducial markers. For DYRK3-inhibition experiments, the cells were treated with 2 µM GSK626616 or 0.0001 % DMSO, 3 hours before fixation with 2.5 % glutaraldehyde (Electron microscopy Sciences) in 0.1M Na+- cacodylate for 1 hour or overnight at room temperature. The cells were then post fixed with 1% OsO4 in 0.1 M Na+-cacodylate, and 1% aqueous uranyl acetate for 1 hour at room temperature. The samples were then dehydrated in an ethanol series and embedded in Epon/ Araldite (Sigma-Aldrich). Ultra thin (70nm) sections were post-stained with lead citrate and examined with a Talos 120 transmission electron microscope at an acceleration voltage of 120 KV using Ceta digital camera and the MAPS software package (Thermo Fisher Scientific, Eindhoven, The Netherlands).

### Correlative Light Electron Microscopy

HeLa cells were grown onto 6 mm sapphire discs described above. 16 hours before the experiment the cells reached about 70 % confluence and were transfected with mCherry-Sec16A IDR using Lipofectamine 2000 as a transfection reagent according to the recommendation of the manufacturer. The cells were then fixed in 0.1M Na+-cacodylate buffer (pH 7.35, prewarmed to 37°C) containing 2% formaldehyde and 0.1% glutaraldehyde (Sigma-Aldrich, Buchs, Switzerland) for 1 h prior light microscopy imaging. Imaging was performed with a spinning disk microscope from Nikon (Eclipse Ti-E) with an enhanced CSU-W1 spinning disk (Microlens-enhanced dual Nipkow disk confocal scanner, wide view type), a Nikon CFI PlanApo 40x oil immersion objective of NA 1.2, and a Photometric Prime BSI sCMOS (2,048 × 2,048 pixels). An overview of the entire slide at low magnification (20x), was acquired using brightfield. The regions of interest for acquisition at 40x magnification were automatically acquire as multiple tiled regions of interest using a custom made focus map. Imaged cells were subsequently processed as described before for cells intended for electron microscopy only. For correlative light and electron microscopy, light microscopy images were loaded into the MAPS software and aligned with the electron microscopy images using three matching points based on fluorescence correlated to the appropriate ultra structure.

### Automated image analysis of immunofluorescence experiments

Image processing, object identification and feature extraction were performed using TissueMAPS (https://github.com/pelkmanslab/TissueMAPS). Every image was corrected for inhomogeneous illumination in TissueMAPS using summary statistics for each stain. Nuclei (primary objects) were segmented using the DAPI signal. Cells (secondary objects) were segmented using the Alexa Fluor 647 NHS Ester signal. Intensity, texture and morphology features were extracted for each segmented object and stain. Multinucleated, mitotic and miss-segmented cells were discarded by supervised machine learning using support-vector machine classifier (SVM). Cells with segmentation masks that extended beyond the field of view (border cells) were also excluded from the analysis. Classification of COPII proteins distribution (punctate or dissolved) and classification of ts045-VSV-G distribution (endoplasmic reticulum-like, Golgi-like, plasma membrane-like or aberrant) were also obtained by supervised machine learning on texture features of the appropriate stainings/ signals. Briefly, an initial training set was obtained by manual assignment of representative cells to each class and the trained classifier was then applied to the assign each cell to a class. Iterative improvement of the training set training accompanied by visual inspection of prediction performance was done to obtain accurate assignment of each cell to its respective class. The segmentation of Sec16A-IDR phospho mutants condensates was obtained using the pixel classifier elastic (http://ilastik.org/ (*91*)) to generate probability maps that were subsequently uploaded in TissueMAPS and used to segment condensates (tertiary objects). Data analysis was performed with custom scripts in R (3.4).

